# Towards a general framework for modeling large-scale biophysical neuronal networks: a full-scale computational model of the rat dentate gyrus

**DOI:** 10.1101/2021.11.02.466940

**Authors:** Ivan Georgiev Raikov, Aaron Milstein, Prannath Moolchand, Gergely G Szabo, Calvin Schneider, Darian Hadjiabadi, Alexandra Chatzikalymniou, Ivan Soltesz

## Abstract

Large-scale computational models of the brain are necessary to accurately represent anatomical and functional variability in neuronal biophysics across brain regions and also to capture and study local and global interactions between neuronal populations on a behaviorally-relevant temporal scale. We present the methodology behind and an initial implementation of a novel open-source computational framework for construction, simulation, and analysis of models consisting of millions of neurons on high-performance computing systems, based on the NEURON and CoreNEURON simulators (Carnevale and Hines, 2006, Kumbhar et al., 2019). This framework uses the HDF5 data format and software library (HDF Group, 2021) and includes a data format for storing morphological, synaptic, and connectivity information of large neuronal network models, and an accompanying open-source software library that provides efficient, scalable parallel storage and MPI-based data movement capabilities. We outline our approaches for constructing detailed large-scale biophysical models with topographical connectivity and input stimuli, and present simulation results obtained with a full-scale model of the dentate gyrus constructed with our framework. The model generates sparse and spatially selective population activity that fits well with in-vivo experimental data. Moreover, our approach is fully general and can be applied to modeling other regions of the hippocampal formation in order to rapidly evaluate specific hypotheses about large-scale neural architectural features.

## 1 Introduction

Large-scale computational models of the brain are necessary to accurately represent anatomical and functional variability in neuronal biophysics across brain regions and also to capture and study local and global interactions between neuronal populations on a behaviorally-relevant temporal scale. To this end, a number of large-scale neural simulation projects aim to facilitate understanding of how the brain’s multi-scale, complex organizational principles comprise cognition and behavior by means of diverse technical approaches (Billeh et al., 2020, Markram et al., 2015). A mechanistic understanding of how individual neural circuits coordinate to generate behavior requires the formulation and systematic exploration of hypotheses about neural computation, which in turns requires the technical ability to rapidly revise and simulate various parameter combinations of diverse biophysical building blocks of neural circuitry on a broad range of spatial and temporal scales.

In the present paper, we describe algorithms and methods that comprise a tentative computational framework for implementation of large-scale neuronal network models and demonstrate its use to construct, simulate, and analyze a full-scale computational model of the rat dentate gyrus (DG). This framework is an embodiment of several core principles that we suggest necessary for large-scale neural modeling efforts:

- Reproducibility
- Flexibility (iterate over hypotheses)
- Scalability (parallel computation, ability to efficiently use computing hardware)

While by no means exhaustive, these principles are necessary for the reproducible and systematic exploration of the parameter space of detailed large-scale neural models.

Our computational modeling framework is structured as follows: 1) an HDF5-based data format for efficient representation of numerical data organized in a hierarchy of cells and cell populations, along with a corresponding library for efficient and scalable parallel data operations; 2) a software library that builds upon our data format and implements algorithms and data structures for construction and analysis of neural volumes, cell morphologies, synaptic distributions, connectivity, and input stimulus patterns, as well as provides tools for running parallel single neuron and network simulations with the NEURON and CoreNEURON simulators (Carnevale and Hines, 2006, Kumbhar et al., 2019); 3) a set of software components that serve to extract subsets of cells from the network and analyze their behavior when provided realistic input spike trains, and optimize their synaptic properties: Network Clamp, Virtual Slice, Microcircuit Clamp.

The next few sections detail these components and illustrate their use in building a full-scale model of the dentate gyrus, a functionally critically important of the hippocampal formation in the temporal lobe of the mammalian brain.

## 2 Data format and scalable parallel I/O format for flexible and efficient representation of neuronal data

The NeuroH5 data format and associated software library is an HDF5-based format for per-cell morphological, synaptic, connectivity and other information of large neuronal network models. The NeuroH5 library provides functions that are intended to assist in the development of scalable parallel algorithms for the construction, simulation, and analysis of network models 1.

Large-scale models require not only high-performance parallel instantiation and simulation, but frequently generate significant amounts of output data and thus require parallel write or append operations in order to store the data, and then parallel read operations for further analysis and visualization of simulation output. Furthermore, during model development it is often desirable to analyze individual cells or subsets of the network model on personal computers. Thus the NeuroH5 data format is designed to provide I/O operations for diverse types of network model and neural simulation result data that scale from personal computers to the largest supercomputers in the world.

The design of the NeuroH5 library is guided by the premise that a core set of general and scalable I/O and data movement utilities can be combined to implement any I/O and data-related operations involved in the construction and analysis of neural network models. Therefore, the NeuroH5 data format is focused on two principal data structures: graph projections, which specify connections between two populations of cells, and cell attributes, which specify numerical attributes associated with individual cells.

Cell attributes have homogeneous data types which cover the most practical cases for numerical computing, that is where the type of the elements is numeric and the precision and representation is efficiently implemented on hardware of most current computer architectures, that is to say 8, 16, 32 and 64 bit integers, either signed or unsigned, and 32 and 64 bit floating point numbers.

The data format is hierarchically organized by neuronal populations, attribute namespaces, and global cell identifiers (gid). Neuronal populations represent a particular kind of neural species, attribute namespaces allow the logical grouping of attributes, and gids identify individual cells within a population. A cell attribute is a homogeneous numerical vector that is contained within a particular namespace and associated with a particular gid in a population. A graph projection is a collection of numerical vectors that specify the source and destination node indices of each edge, where each node index corresponds to a gid.

Correspondingly, the NeuroH5 library provides a set of scalable routines for parallel reading, writing and movement of data, in order to assist model developers in efficient parallel operations for building, simulation, and analysis of the model. Specifically, the following general operations:

1. parallel read, two-phase scatter/read, selection read of cell attributes, graph edges, and node attributes;
2. parallel write, two-phase gather/append of cell attributes, graph edges, and node attributes;
3. broadcast of cell attributes and graph edges
4. parallel Python generator which permits the sequential partial parallel two-phase scatter/read operations of data sets.

The NeuroH5 software library is written in C++ and uses MPI (Message Passing Interface) 2.0 (Forum, 2015), the HDF5 library (HDF Group, 2021), and the Cereal C++ library for serialization (Grant and Voorhies, 2017). In addition, it provides a Python interface that represents and manipulates the data as NumPy arrays (Harris et al., 2020). Efficient, parallel I/O is often a bottleneck in HPC applications, especially those that read and write large amounts of data from and to parallel file systems. MPI-IO (Thakur et al., 1999) (part of MPI-2) serves as the foundation upon which higher-level parallel I/O libraries such as parallel HDF5 (Chaarawi and Koziol, 2012) are built. The NeuroH5 data format and software library is based on the HDF5 format and uses the MPI-specific parallel HDF5 functionality to support efficient block-structured, two-phase I/O mechanisms (del Rosario et al., 1993).

Next, we discuss the data structures used to organize NeuroH5 data sets and the various mechanisms for distributing the data once it is read, or for collecting it for writing.

### 2.1 Data Structures

NeuroH5 data structures were designed with memory scalability and efficient parallel I/O in mind. HDF5 uses a file directory-like structure that allows data to be organized in a nested hierarchical structure, and accordingly the NeuroH5 uses a hierarchical format that is organized by population name, which allows data for cells that belong to the same population to be grouped together, and attribute namespace, which allows related data attributes to be grouped together (Figure 2). A set of tables defined as structured data types in HDF5 provide information about the valid population name and indices of cells within each population.

**Figure 1:**
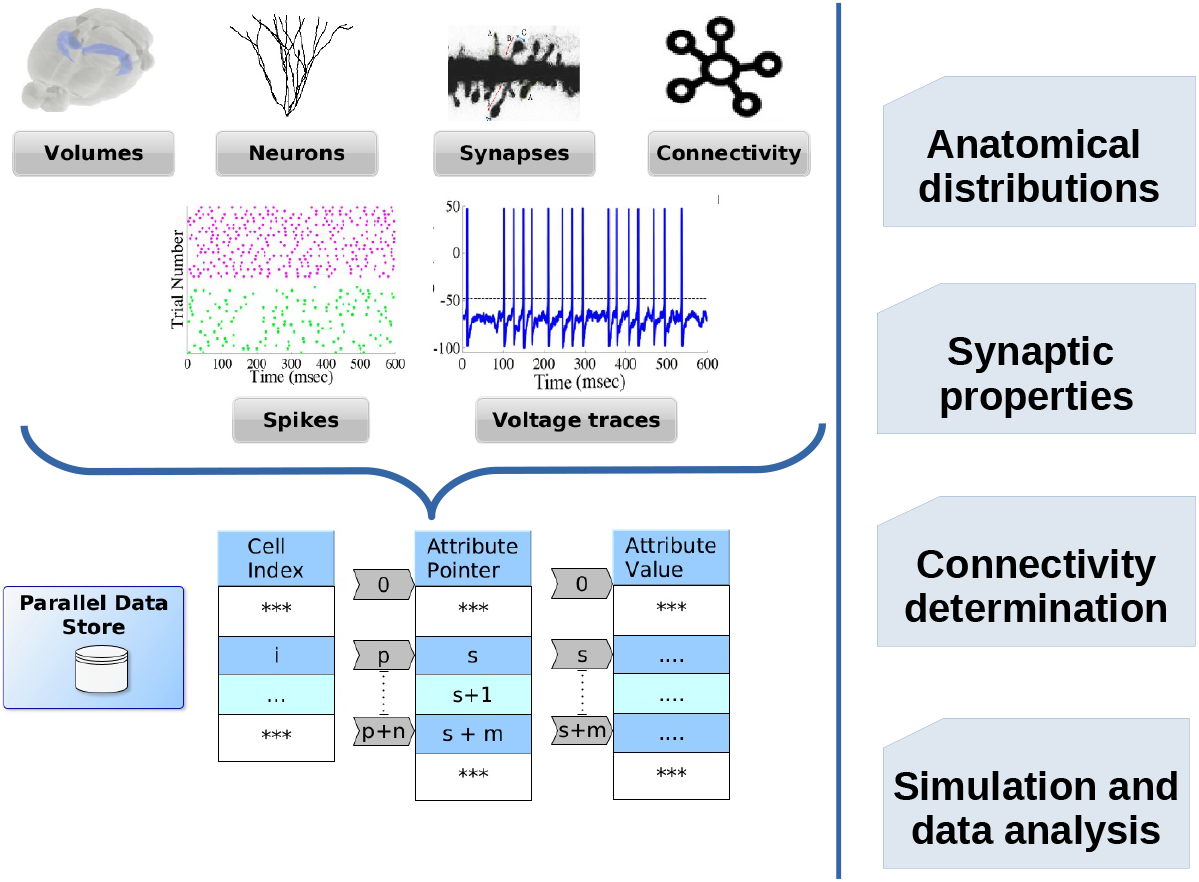
Modeling workflow for large-scale biophysical neuronal network models supported by the NeuroH5 data format and software library. NeuroH5 offers data structures that allow diverse types of morphological, synaptic, and connectivity neuron model data, as well as spike and state variables recordings to be efficiently read and written in parallel. Common operations involved in model construction, instantiation, and analysis can use the uniform data interface provided by NeuroH5.

**Figure 2:**
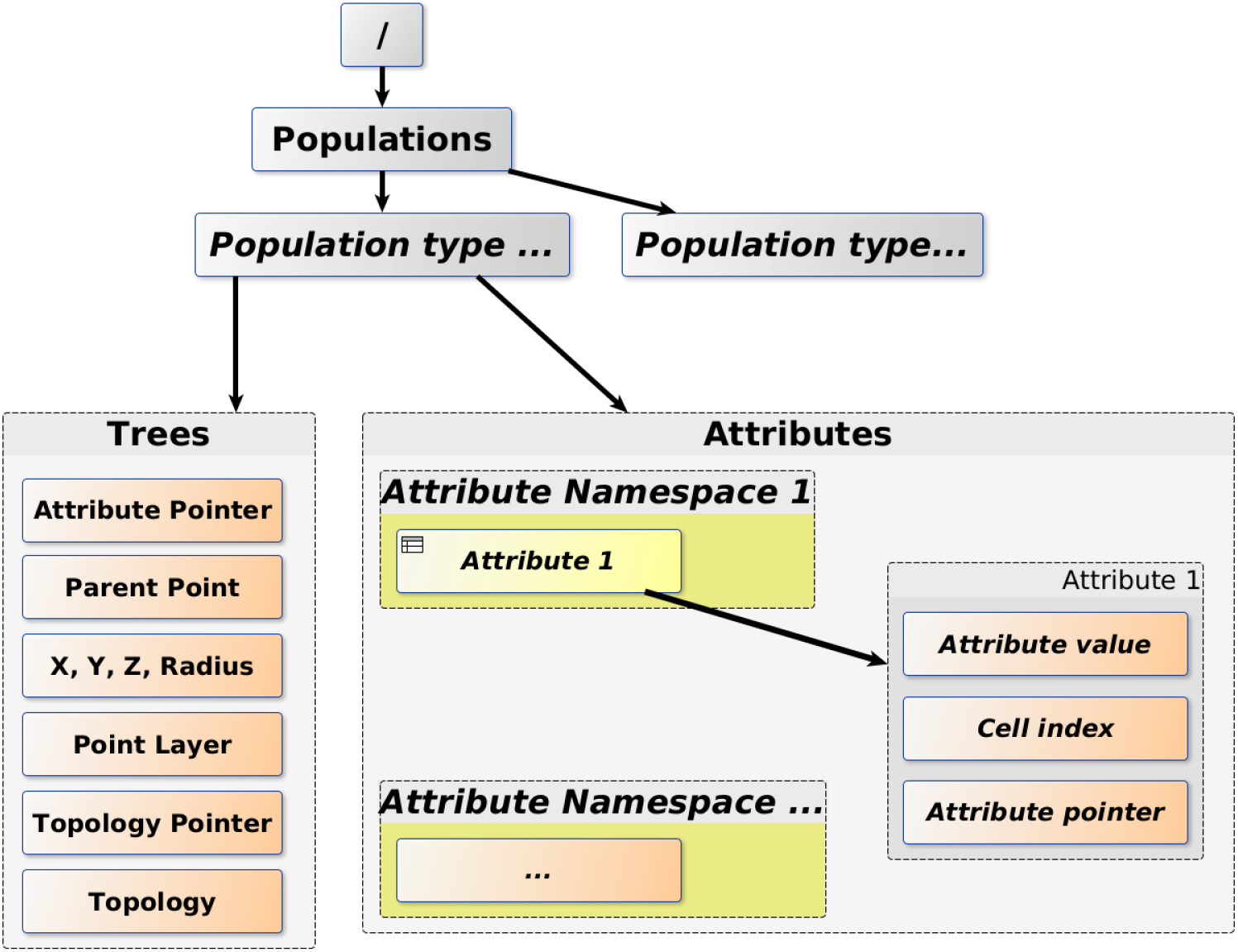
An illustration of the hierarchical structure of NeuroH5. NeuroH5 cell attributes are organized by population and attribute namespace. Within a population, there are multiple namespaces, which in turn contain multiple attributes. In this example, the “Trees” namespace contains the cell attributes that describe the SWC morphology format. A cell attribute is described by three datasets, attribute value, cell index, attribute pointer. See text for further details.

The data for an individual cell attribute is described with three HDF5 datasets: cell index, attribute pointer, attribute value. Graph connectivity data is described in a format inspired by sparse graph representations that we have termed Destination Block Sparse.

A cell attribute is described with three datasets (Figure 2):

1. The `Attribute Pointer` dataset contains the data offset that indicates the starting position of the attribute data for the corresponding cell. In Figure 2, the `Cell Index` dataset contains index *i* at position *p*, and the `Attribute Pointer` dataset contains offset *s* at position *p*. This means that the data for cell index *i* begins at position *s* in the `Attribute Value` dataset. In addition, `Attribute Pointer` dataset contains offset *s* + *k* at position *p* +1. This means that the data for the next cell in this attribute set begins at position *s* + *k* in the `Attribute Value` dataset, where *k* is the number of elements associated with cell index *i*.
2. The `Attribute Value` dataset contains the attribute values for all cells of a given population that are associated with this attribute. As described above, the `Attribute Pointer` dataset contains each offset *s* that points to the beginning of the attribute data for a given cell index *i*.

Figure 3 provides an example of model cell data encoded in the NeuroH5 format. Specifically, the widely used SWC morphology representation (SWC, 2021) is represented as one attribute for each of the SWC data fields, XYZ coordinates, radius, point connectivity, type, as well as additional attributes not part of the original SWC format, such as layer and section grouping.

**Figure 3:**
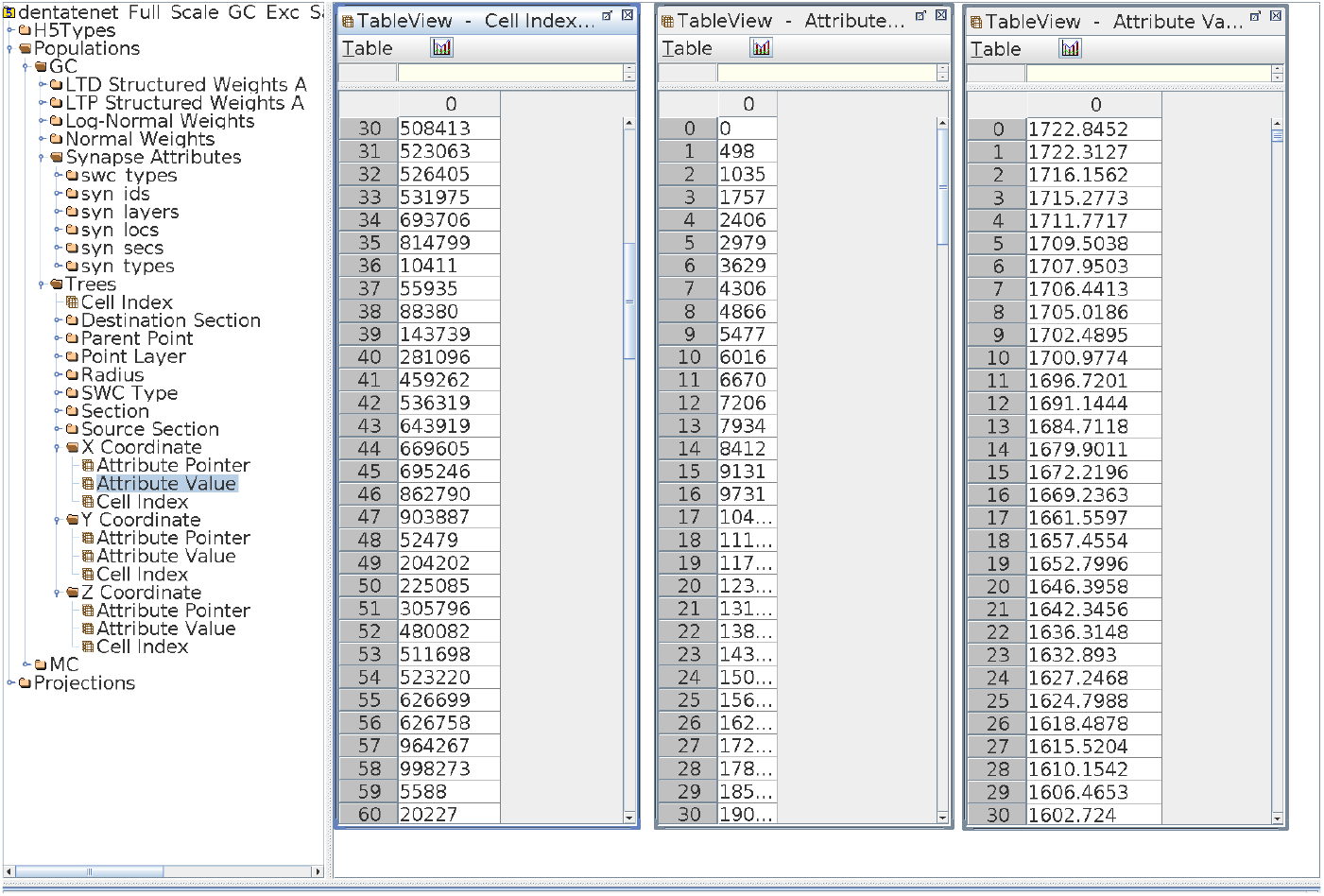
Example NeuroH5 data sets. Leftmost panel: Contents of a NeuroH5 HDF5 file with dentate granule cell morphologies and synaptic attributes. Right panels: the contents of the NeuroH5 datasets that describe the X coordinates of all morphological points of the population of granule cells.

This simple data architecture has proven sufficiently flexible to express a broad range of cell attributes, as we will see further. The index and pointer data structure permit the data to be split by cell index efficiently, while the single, contiguous value attribute dataset permits the data to be divided in regions that can be read efficiently in parallel.

Next, we describe the connectivity data format of NeuroH5. Connectivity graphs are organized in hierarchical structure organized by destination and source populations. Each projection is described in terms of source and destination nodes (which are commonly also called vertices Each node index corresponds to a cell index. By default, the Destination Block Sparse format represents connectivity as a destination node index associated with a set of source node indices, but the format supports source node index-based address as well. We define a block as a subset of the connectivity graph that contains a contiguous set of destination node indices.

Figure 4 shows an example directed graph and its representation in the NeuroH5 Destination Block Sparse format. The example graph contains 10 nodes and 14 directed edges. For example, there is a directed edge from node 12 to node 13. Nodes 10 and 11 have no incoming edges.

**Figure 4:**
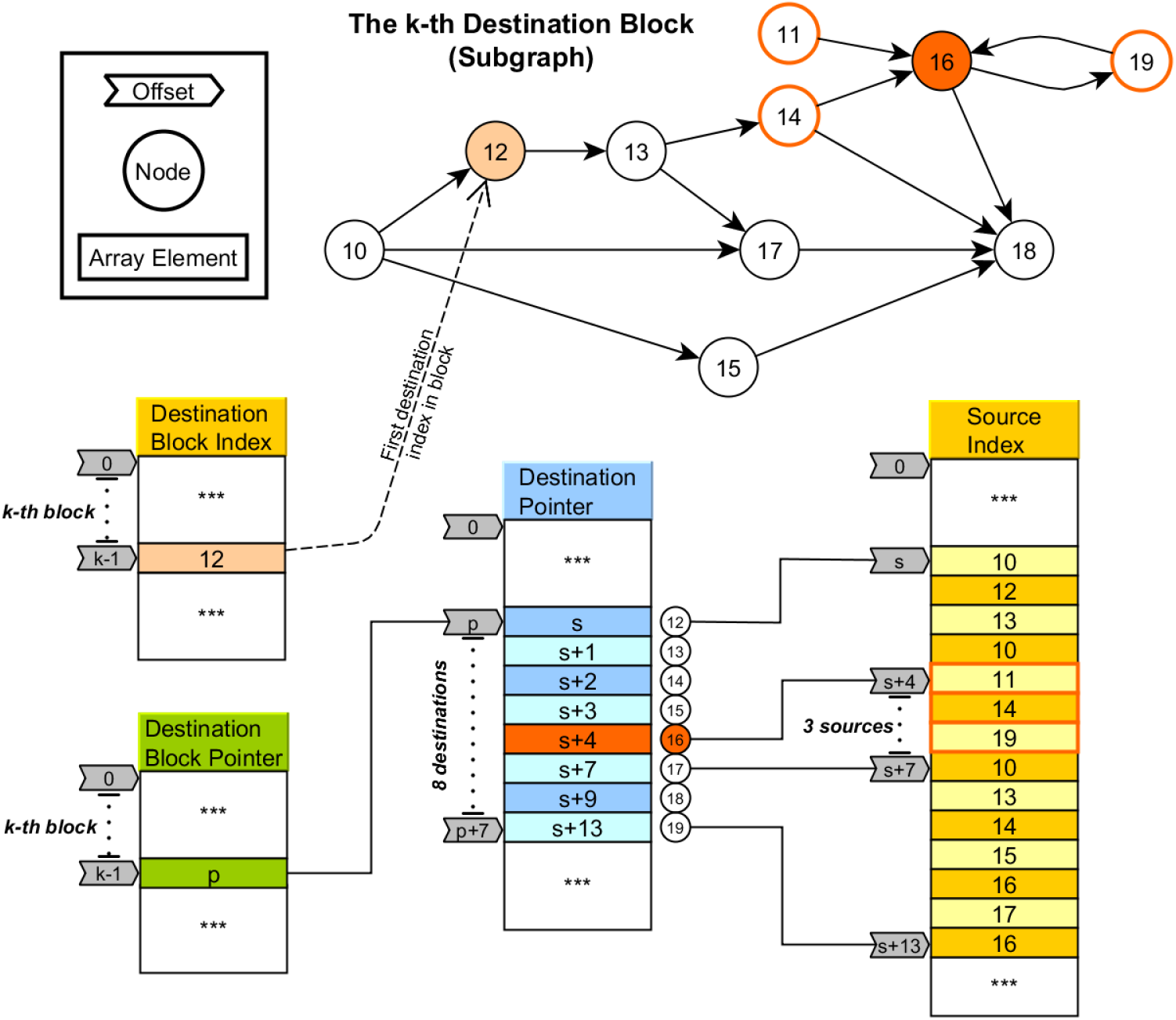
Example of connectivity structure encoded in NeuroH5 Destination Block Sparse format.

Graph connectivity is then described by four datasets (Figure 4):

1. The `Destination Block Index` dataset contains the starting node index of a contiguous set of destination node indices, where a node index corresponds to the relative cell index. Each such contiguous set of destination node indices is termed a *block*. In the example in Figure 4, the subgraph depicted at the top is described by a block where the first node index is 12.
2. The `Destination Block Pointer` dataset contains the offset in `Destination Pointer` dataset corresponding to each destination node in the block. In the example in Figure 4, block *k* contains nodes 12… 19, and the starting offset for block *k* can be found at position *p* in the `Destination Block Pointer` dataset.
3. The `Destination Pointer` dataset contains the offset in `Source Index` dataset corresponding to beginning of the vector containing source node indices for each destination node in the block. In the example in Figure 4, block *k* contains nodes 12… 19, the starting offset for block *k* can be found at position *p* in the `Destination Pointer` dataset, and each element in positions *p*… *p* +7 specifies an offset *s* that indicates the starting position of the source indices for each node in the block.
4. The `Source Index` dataset contains the indices of the source nodes corresponding to each destination node in the block. Figure 4, block *k* contains nodes 12… 19, the starting offset for block *k* can be found at position *p* in the `Destination Pointer` dataset, and each element in positions *p*… *p* + 7 accordingly specifies offsets *s*… *s* + 13 that indicates the elements of the `Source Index` dataset that contain the corresponding source node indices. Thus, the source indices for destination node 16 can be found at position *p* + 4 in `Destination Pointer`, which contains the value *s* + 4, and in turn position *s* + 4 in `Source Index` points to the beginning of a vector containing the elements 11, 14, 19, i.e. the source indices of node 16 in the example graph.

The Destination Block Sparse format was inspired by the Compressed Sparse Column (CSC) and Compressed Sparse Row (CSR) graph formats, which originated in high-performance scientific computing as a way to represent sparse matrices that contain mostly zeros. Those formats allowed the efficient packing of row or column indices of non-zero into dense arrays, which allows for a compact, contiguous memory layout, which offers significant space saving and efficient memory access. The trade-off is flexibility: graph analysis and manipulation of a destination-oriented representation will be inefficient if a source-based graph traversal is required, and vice versa. Our design choice of default destination-based format was dictated by the observation that neuronal network models typically represent connections from the perspective of the post-synaptic neuron, which corresponds to the destination node in the graph. The NeuroH5 format alternatively supports a source-based connectivity representation format. In either case, the contiguous layout of the connectivity dataset again permits efficient indexing and partitioning for parallel I/O operations.

### 2.2 Operational Architecture

The data access patterns provided by the NeuroH5 library are either reading/writing contiguous blocks of data or selection reads that access only the data for specific non-contiguous cell indices. The principal parallel I/O operation is based on a two-phase approach where a set of designated I/O processes consolidate numerous data requests into few large contiguous reads. Data are gathered from and distributed to all processes via the MPI collective operation `MPI_Alltoallv`.

The basic architecture of the NeuroH5 library is depicted in Figure 5. The user specifies the I/O group size and basic parameters of the I/O operation (file name, read/write, contiguous/selection, offsets, data dimensions etc.). All I/O to and from disk is performed in parallel via HDF5 and MPI I/O. Algorithm 1 illustrates the basic sequence of steps that comprise the block scatter/read operation.

**Figure 5:**
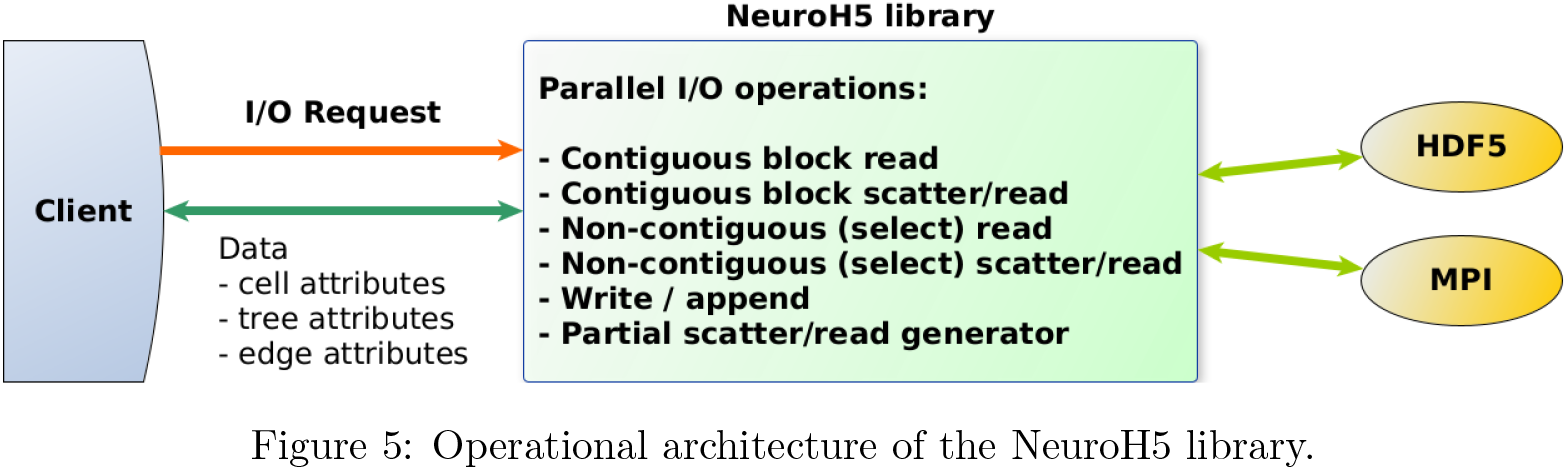
Operational architecture of the NeuroH5 library.

**Listing 1:**
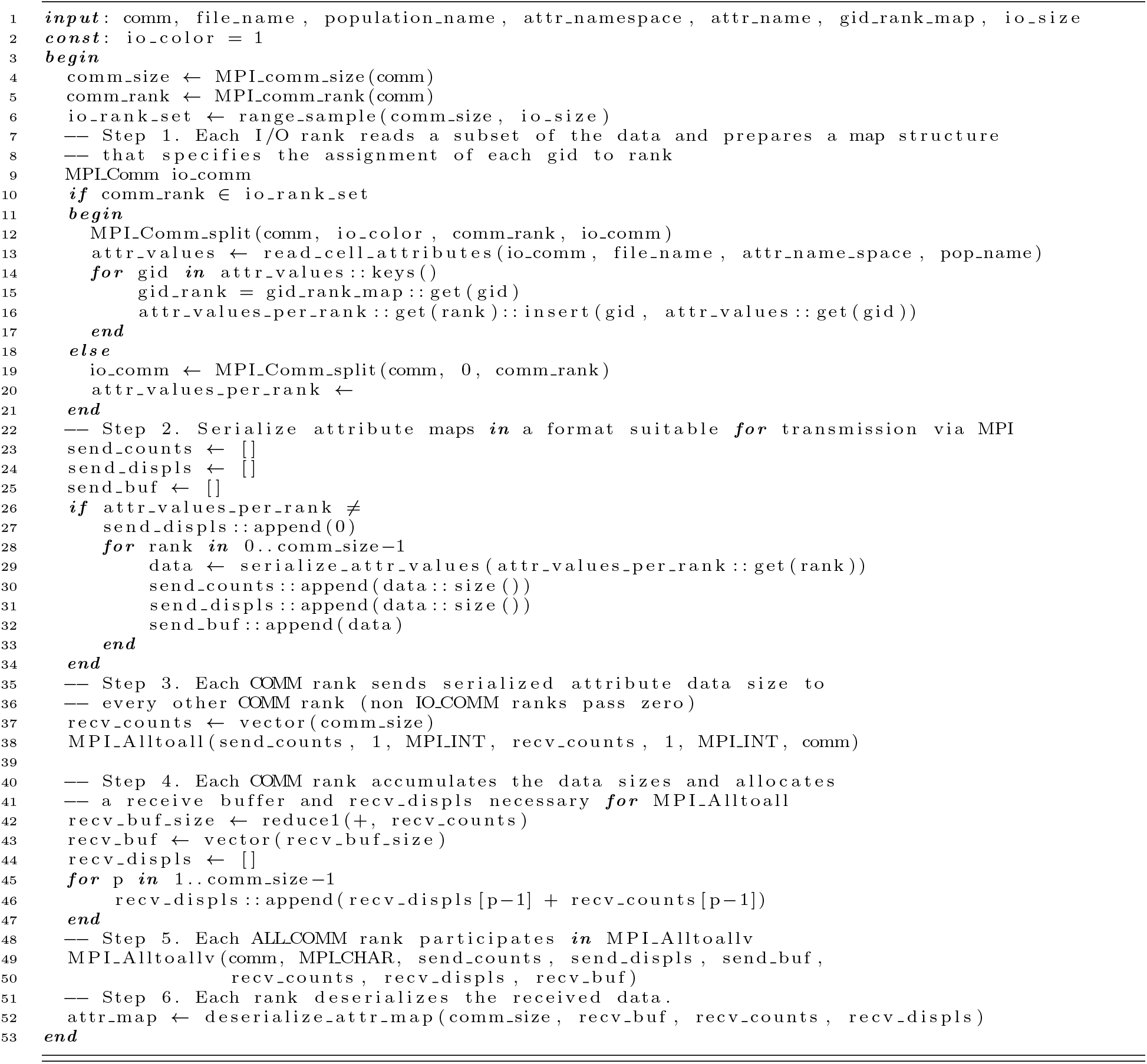
NeuroH5 two-phase block scatter/read operation for cell attributes

## 3 Reproducible and detailed representation of neural geometry, connectivity, and inputs

Detailed three-dimensional geometric representations of neuroanatomical volume and neuronal morphology are essential for computational modeling of neuronal networks. The three-dimensional geometry of the neural volume is a critical determining factor of the shape of neuron layers, the neurons’ morphological structure and spatial projection patterns of their axons. We have developed a set of methodologies, based on parametric geometric surfaces, to derive synaptic distributions, network connectivity and conduction delays from anatomical reconstruction data, as well the dendritic and axonal projection distributions. A mapping technique enables the projection of the 3D positions in the volume to a two-dimensional map where connectivity can be computed based on the arc-distances between somata in the respective layers. In section 5.1 we will demonstrate how this methodology can reveal complex properties of the synaptic connectivity patterns.

### 3.1 Neural volume geometry

Fitting and approximating neuroanatomical reconstruction data efficiently, and yet with sufficient resolution, are connected with fundamental research problems in computational geometry with a long history of study. While it would be ideal to directly use three-dimensional point data without any loss of information, the limitations of memory and compute time dictate that a parsimonious mode of representation is crucial.

Among the well-established form of three-dimensional geometric data are those based on parametric curves, which specify point coordinates as functions of one or more independent variables. They provide the ability for arbitrarily smooth surfaces, which makes them especially suitable for computer-aided design. Therefore, the conversion of neuroanatomical reconstructions from point cloud data to parametric representations is a fundamental step in the creation of highly realistic computational models of the brain. However, generating accurate geometric data with parametric surfaces based on experimental observations often requires approximation and a general and accurate fitting solution is difficult to obtain.

In the present study, we used a previously developed parametric volume representation of the dentate gyrus (DG) (Schneider et al., 2014). Smoothed surfaces for the boundaries of the DG granule cell layer (GCL) and molecular layer (ML) were obtained from a high-resolution, 3D serial reconstruction of the rat hippocampus (Ropireddy et al., 2012) (Figure 6. This in turn provided realistic geometric context based to provide constraints for an algorithm for generation of realistic dendritic morphology (Schneider et al., 2012, 2014). In brief, a set of points are distributed within a three-dimensional cone approximating the shape of the granule cell dendritic field and connected for each individual cell via an optimal wiring algorithm (Cuntz et al., 2012, 2010). Spatial jitter and diameter mapping are then added to reproduce the tortuosity and quadratic diameter tapering, respectively, of real dendrites.

**Figure 6:**
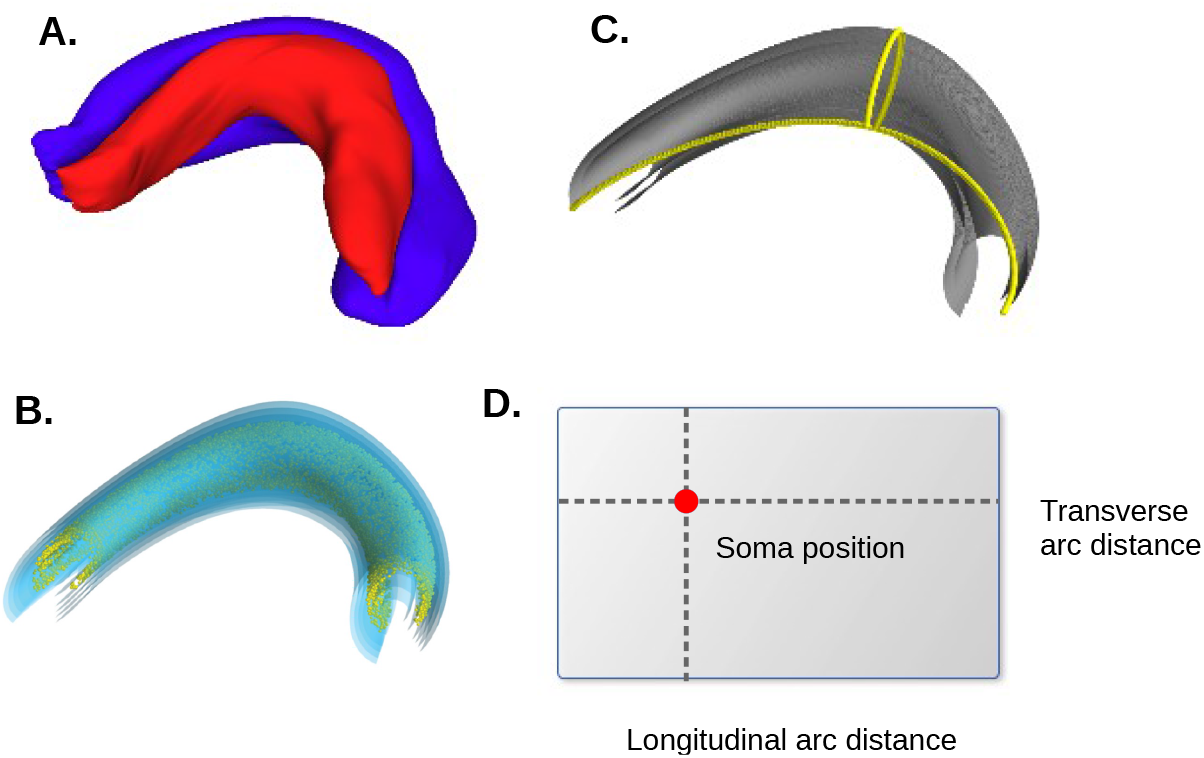
Parametric volume representation of neural geometry. A. Rendering of the hippocampal volume reconstructed by Ropireddy et al. (Ropireddy et al., 2012) B. Parametric volume fit to the reconstructed dentate gyrus volume, with cell somata distributed within the volume. C. A set of points distributed along the longitudinal and transverse axes of the parametric volume in order to calculate arc distances within the volume. D. A representation of the projection to 2D anatomical space used to calculate distances between cell positions in the dentate gyrus model.

The use of a parametric surface based on experimental data has permitted the linking of neural volume geometry with the anatomy of individual dendrites and topographical connectivity between neurons. Our framework makes it possible to seamlessly link neuroanatomical features with gradients of morphological, electrophysiological and connectivity properties, and answer detailed questions of how the anatomically constrained distributions of these properties affect the functional properties of the region under study.

### 3.2 Layer-based topographical connectivity in 3D

Once a parametric surface representation of the anatomical volume is created, neuronal somata can be distributed to approximate the layer and extents of experimentally observed somata distributions. Our framework provides a method to disperse cell somata within a domain bounded by parameter surface coordinates that reflect the appropriate layer constraints. The dispersion is analogous to electrostatic repulsion, where neighboring nodes exert a repulsive force on each other. Each node is moved in the direction of its net repulsive force with a step size proportional to the distance to its nearest neighbor.

Next, synaptic locations are determined based on anatomical data about neurite location and layer-specific synapse density. Spacing between synapse locations is determined by a Poisson distribution with parameters derived from the synapse density and compartment length at each segment of the cell morphology. The use of a parametric surface allow the layer of a particular point on the dendritic tree to be determined at an arbitrarily high resolution.

Once soma positions and synaptic locations are defined, connectivity patterns in our framework are expressed in terms of parameters that denote the anatomical layer, neurite location, and pre-synaptic source of the target synapses. Network connectivity is defined solely on the level of populations of neurons and not for single neurons. Using a simple mapping technique (see below), somata locations in different layers are projected to a 2-D arc-distance space in order to create layer-dependent probability distributions of neuronal connectivity that are based on experimentally determined axonal extents. These distributions are then used to determine connections and compute distances between neurons which in turn is used to compute the transmission delay. This framework allows the easy expression of complex and realistic anatomical connection patterns between populations of neurons.

Using probability functions reflects the assumption that all neurons that belong to the same neuron type have the same genetic code and emerged through cell proliferation processes that gave rise to qualitatively similar connectivity patterns.

A consequence of the use of parametric surfaces is that any point in the volume can be described by its XYZ-coordinates in Euclidean space and in parametric surface coordinates, sometimes referred to as UV-coordinates in the computational geometry literature. This property allows the description of anatomical properties either along Euclidean space axes or along anatomical axes which are not necessarily aligned with Euclidean axes, and therefore give rise to very different connectivity patterns.

The parametric surface approach allows a more accurate conversion of anatomical observations to complex non-linear patterns of axonal projections and the location and dendritic tree orientations of neurons. This is critical for large-scale hippocampal networks, as projection patterns within the hippocampal layers (dentate gyrus, CA3, CA1) follow local axes such as proximal-distal, septal-temporal (Andersen et al., 2006), which are determined by the shape of the hippocampal volume, and therefore connectivity determined solely by Euclidean distance will be highly distorted.

Our parametric surface approach allows complex connectivity patterns based on anatomical distances to be efficiently represented and connectivity distributions to be rapidly generated.

### 3.3 Topographical representation of spatial input gradients

The major cortical inputs to the hippocampus are provided by neurons in layer II of the entorhinal cortex (EC) (Tamamaki and Nojyo, 1993). EC is functionally organized into two components, the lateral EC (LEC) and medial EC (MEC) (Witter et al., 2017). MEC and LEC contain topographically organized cells that encode representations of position, movement direction and velocity (Sargolini et al., 2006). In particular MEC, contains a significant population of grid cells, which are place-modulated neuron whose receptive fields are organized in a hexagonal grid pattern (Hafting et al., 2005). Stensola et al. (Stensola et al., 2012) found that grid cells were organized in 4-5 modules, extending over 50% of the dorsoventral MEC and by extrapolation hypothesized that around 10 modules exist in the entire MEC. Grid cell spacing were distributed in discrete logarithmic increments: 40, 50, 70, 100 *cm* and organization of grid orientation and scale was comodular.

Based on the results by Stensola and colleagues (Stensola et al., 2012), we assumed that extrinsic inputs from MEC, LEC, and CA3 have grid spacing and/or spatial field widths that are topographically organized septo-temporally. Input cells that are designated to be grid cells are assigned to one of ten discrete modules with distinct grid spacing and field width sampled from Gaussian distributions according to septo-temporal position. The grid spacing across modules increases exponentially from 40 *cm* to 8 *m*. Input cells that are designated to be place cells have place fields that result from sampling input from multiple discrete modules, and therefore have field widths that vary continuously with septo-temporal position, rather than clustering into discrete modules. Furthermore, we create populations of “proxy input cells” that correspond to GC and MC populations with idealized distributed of spatial selectivity features. These “proxy input cells” are used to provide generate spike trains that are outside the target volume during Microcircuit Clamp or Network Clamp simulations, described next.

## 4 Network Clamp and Microcircuit Clamp

One of the significant challenges of large-scale neuronal network models is that their high-dimensional parameter space is often difficult to constrain without detailed data about synaptic and cellular biophysical properties. In order to allow the study of model cell behavior in the context of large-scale network dynamics, we have developed two related approaches termed Network Clamp and Microcircuit Clamp. The Network Clamp concept was first developed for our previous data-driven, full-scale model of the CA1 region (Bezaire et al., 2016b) and was first described by Bezaire et al. (Bezaire et al., 2016a).

In the Network Clamp approach, a single target cell is extracted from the full-scale network, along with all of its synaptic connections and the corresponding cell type-specific presynaptic activity patterns, so that all state variables of the target cell can be analyzed and recorded while the cell receives realistic input from the network. Network Clamp also provides the ability to generate arbitrary network inputs on any synapse in order to test the cell behavior in contexts other than that of the model network. This dramatically reduces the computational resources required for simulation without losing the strong biological realism of the biophysical full-scale model.

The Network Clamp approach provides a convenient platform for synaptic parameter search and optimization. We have developed an optimization methodology, based on Network Clamp and global optimization of Lipschitz functions (Malherbe and Vayatis, 2017), wherein the synaptic weights can be optimized to match a target firing rate when the cell is provided inputs with experimentally constrained firing rates.

The closely related Microcircuit Clamp approach involves the extraction of a virtual slice of the full-scale network and performing simulations and parameter optimization over the cells contained in the slice volume, while providing appropriate spike trains in place of the neurons outside of the volume. The Microcircuit Clamp likewise permit analysis and optimization over the synaptic parameters of the network in order to match target constraints that are possibly derived from experimental data.

## 5 Results

### 5.1 A full-scale, data-driven, biophysical neural network model of the rat dentate gyrus with realistic spatial input

The hippocampus provides the basis for spatial navigation and episodic memory in the brain, storing and recalling events experienced in the past and linking them with their spatio-temporal context (Buzsáki and Moser, 2013, O’Keefe and Nadel, 1978). The dentate gyrus (DG) is one of the major subfields of the hippocampus, with excitatory input arriving from Layer II of the entorhinal cortex (EC). The principal cells of the dentate gyrus are the granule cells (GCs), which are glutamatergic projection neurons that constitute 99% of the cell population in the DG and are thought to have a central role in memory formation.

A striking property of the dentate gyrus is that in an awake behaving animal, only a small fraction (1—10%) of the GC population is concurrently activated by the EC inputs, yet this is sufficient to imprint memories onto the downstream CA3 region. Since only a small set of narrowly tuned GCs appear to be involved in each memory representation, this phenomenon has been called “sparse population coding” (Olshausen and Field, 2004), and is thought to be the means to achieve robust yet parsimonious encoding of information in the nervous system. A major advantage of sparse population coding is the ability to store non-overlapping representations of similar inputs and correctly retrieve them given a partial or noisy cue stimulus. Theoretical models refer to those processes as pattern separation and pattern completion, respectively.

The cellular and circuit characteristics of the dentate gyrus combine uniquely to constrain the number of active GCs in order to maintain sparse population activity. GCs have a high action potential threshold, which is a consequence of their particularly negative resting membrane potential and relatively low input resistance and furthermore, their dendrites are passive and leaky, thereby strongly attenuating synaptic inputs along the dendritic tree. Similarly to the rest of the hippocampal formation, GCs receive strong inhibition from local GABAergic interneurons (INs). However, a very conspicuous feature of the dentate circuitry is that GCs also receive direct excitatory glutamatergic input from mossy cells (MCs). Mossy cells also receive backprojections from CA3c, thus forming a feedback loop from CA3 to DG (Scharfman, 2007). Although the functional role of mossy cells is a topic of active research, they have been shown to play an active role in spatial memory (Bui et al., 2018), and may support pattern separation by recruiting inhibition to control granule cell sparseness (Myers and Scharfman, 2011).

Taken together, these properties of the GCs and of the DG circuitry lead to the hypothesis that overlapping EC input patterns are separated through distinct populations of GCs that are strongly modulated by local interactions of MCs and INs, as well as non-uniform distributions of excitatory synaptic weights shaped by long-term potentiation (Bromer et al., 2018).

In the present study, we constructed a full-scale, data-driven computational model of the rat DG to investigate the activity of GC, MC and different IN populations in the encoding of realistic spatial trajectory input (Figure 7). By applying entorhinal stimuli that mimicked spatial receptive fields, we find that through the distinct inhibitory circuits excitatory, and non-uniform mechanisms, the principal neurons in DG provide sparse and selective activity in response to largely overlapping input patterns.

**Figure 7:**
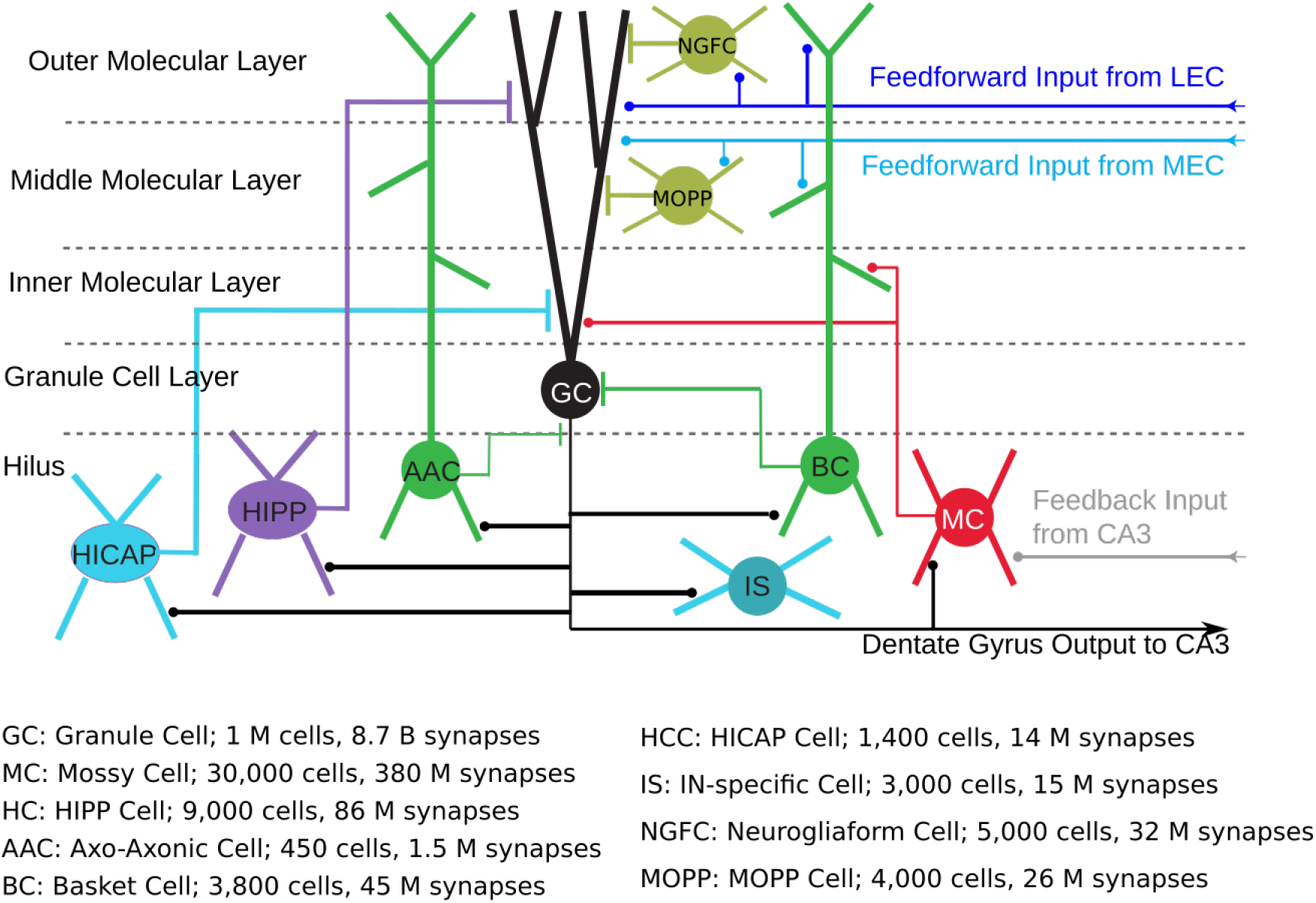
Schematic of the connectivity structure of full-scale dentate gyrus model. Connections between interneurons are omitted for clarity.

### 5.2 Parametric Representation of Neural Volume

The three-dimensional structure of the model rat dentate gyrus and laminar divisions were taken from a previous study that matched several experimental volumetric and width estimates (Schneider et al., 2014) (Figure 6). The parametric equations that generate XYZ volume coordinates were the following:

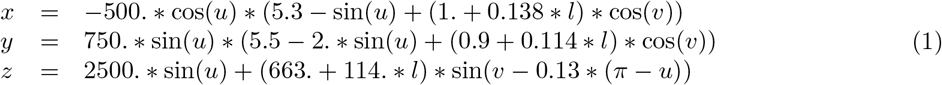

where:

- *v* defined the transverse axis (the C-shape) and ranged from —0.23*π* to 1.425*π*
- *u* defined the septotemporal (or longitudinal) axis and ranged from 0.0*π* to 0.98*π* for the hilus and granule cell layer (GCL) and —0.016*π* to 1.01*π* for the molecular layer
- *l* defined the layer and was −1.95 for granule cell layer (GCL), 1 for inner molecular layer (IML), 2 for middle molecular layer (MML), and 3 for outer molecular layer (OML).

The hilus layer was created using −3.95 for the *l* parameter for the inner boundary to generate a 2.52 *mm*^3^ volume below the granule cell layer, calculated using the experimental 0.6 GCL to hilus volumetric ratio and their combined volume of 6.30 *mm*^3^ (Ropireddy et al., 2012). The total transverse arc length of the OML layer was 5.4 *mm* and the total longitudinal arc length of the OML layer was 11.7 *mm*.

### 5.3 Distribution of Cell Types

The number of neurons and laminar distribution for each cell type in the model are summarized in Table 1. These have been updated from our previous models (Dyhrfjeld-Johnsen et al., 2007, Morgan and Soltesz, 2008, Santhakumar et al., 2005) based on recent immunohistochemical studies. The total number of granule cells in the rat dentate gyrus is estimated to be 1 million (Freund and Buzsáki, 1998, Patton and McNaughton, 1995). Buckmaster and Jongen-Relo (Buckmaster and Dudek, 1999) estimated the number of cells that do not express GAD67-mRNA (a marker for GABAergic interneurons) in the hilus to be 30,000, which are thought to primarily consist of mossy cells. Huusko et al. (Huusko et al., 2015) recently estimated the laminar distribution and total number of neurons for several neurochemical markers throughout the hippocampus. They reported the total number of PV+ neurons in the rat dentate gyrus to be approximately 4,250, and the laminar distribution was approximately 1900, 1900, and 450 for the hilus, granule cell layer (GCL), and molecular layer (ML), respectively. Parvalbumin-positive (PV+) neurons are comprised of basket cells and axo-axonic cells in the dentate gyrus (Ribak et al., 1990, Soriano et al., 1990), and 5 out of 83 ( 6%) recorded PV+ cells in a recent dentate gyrus study were axo-axonic cells (Hu et al., 2010), which is slightly less than an earlier 10%-15% estimate of axo-axonic cell abundance within CA1 (Baude et al., 2007). The number of axo-axonic cells was chosen as the midpoint between these two approximations ( 10.5%), resulting in 3800 PV+ basket cells and 450 axo-axonic cells. These cell types were then distributed based on the previously mentioned laminar ratio.

**Table 1:** Layer distribution of cell somata and external inputs in full-scale model of the dentate gyrus.

|  | Hilus | GCL | IML | MML | OML |
| --- | --- | --- | --- | --- | --- |
| Cell or input type |  |  |  |  |  |
| Granule | 0 | 1,000,000 | 0 | 0 | 0 |
| Mossy | 30,000 | 0 | 0 | 0 | 0 |
| Axo-Axonic | 450 | 0 | 0 | 0 | 0 |
| Basket | 1900 | 1900 | 0 | 0 | 0 |
| HIPP | 9,000 | 0 | 0 | 0 | 0 |
| HICAP | 1,400 | 0 | 0 | 0 | 0 |
| IS | 3,000 | 0 | 0 | 0 | 0 |
| MOPP | 0 | 0 | 2000 | 1000 | 1000 |
| NGF | 0 | 0 | 0 | 2500 | 2500 |
| CL Mossy | 0 | 0 | 30,000 | 0 | 0 |
| MPP | 0 | 0 | 0 | 38,000 | 0 |
| LPP | 0 | 0 | 0 | 0 | 34,000 |
| CA3c | 67,000 | 0 | 0 | 0 | 0 |

HIPP (hilar perforant pathway-associated) cells are thought comprise the somatostatin-positive (SOM+) population in the hilus (Freund and Buzsáki, 1998, Katona et al., 1999) which is approximately 9,000 cells (Buckmaster and Dudek, 1999, Huusko et al., 2015). There are also an estimated 1,400 cholecystokinin-positive (CCK+) cells in the rat dentate gyrus (Buckmaster and Dudek, 1997, Huusko et al., 2015), of which approximately 1150 and 250 cells are located in the hilus and GCL, respectively (Huusko et al., 2015). The CCK+ population consists of two reported cell types, HICAP (hilar commissural-associational pathway-related) cells and CCK+ basket cells. Owing to lack of information about CCK+ basket cells, these two cell types were combined. The number of GAD67-mRNA positive cells in the molecular layer is estimated to be up to 10,000 (Buckmaster and Dudek, 1999), and approximately 62% to 80% of these cells express neuronal nitric oxide synthase (nNOS) (Jinno et al., 1999, Liang et al., 2013). Neurogliaform cells in the rat dentate gyrus are located in the outer two-thirds of the molecular layer and express nNOS (Armstrong et al., 2011). Assuming an equal distribution of interneurons throughout the molecular layer, the number of neurogliaform cells was estimated as 5,000, and the number of MOPP cells (molecular layer interneurons with axons in perforant-path termination zone) with somata located in the inner and middle molecular layers was estimated to be 4,000.

As mossy cells have extensive commissural projections (Frotscher et al., 1991) with almost all cells projecting bilaterally (Deller and Frotscher, 1997), a population of contralateral mossy cell inputs was created, with the same population size as the mossy cell population and distributed exclusively in the inner molecular layer.

In order to represent inputs to the dentate network from Layer II medial and lateral entorhinal cortex (MEC and LEC), we created distributions of EC axon entry points in the middle and outer molecular layer, respectively for MEC and LEC inputs. It has previously been estimated that 67% of Layer II EC cells project to the dentate gyrus, and therefore the number of entry points corresponded to 38,860 MEC inputs (67% of 58,000 cells) and 30,820 LEC inputs (67% of 46,000 cells) (Gatome et al., 2010, Mulders et al., 1997). Furthermore, in order to reflect the existence of an excitatory back-projection from CA3c to the basket cells and mossy cells in the dentate gyrus that has long been reported (Kneisler and Dingledine, 1995, Scharfman, 1994), we have added a population of CA3c inputs equal to the estimated number of pyramidal cells in CA3c (67,000).

The septotemporal and layer distributions for all cell types are summarized in Table 1. Granule cell somata were modeled as randomly placed non-overlapping ellipsoids of 10.3 *μm* width and 18.6 *μm* length, the average values reported in a quantitative study of 48 granule cells (Claiborne et al., 1990). All cell somata were distributed without overlap within the bounds of the volume defined by their respective layers by means of the electrostatic repulsion model outlined in section 3.

### 5.4 Neuron Models

One million unique granule cell dendritic morphologies were generated inside of a reconstructed three-dimensional structure based on our recently established methodology (Schneider et al., 2014). In brief, target points are selected within a three-dimensional cone (the shape of the granule cell dendritic field) and connected for each individual cell via an optimal wiring algorithm (Cuntz et al., 2012, 2010). Spatial jitter and diameter mapping are then added to reproduce the tortuosity and quadratic diameter tapering, respectively, of real dendrites. The morphologies were populated with the mechanisms from the compartmental granule cell due to Aradi and Holmes (Aradi and Holmes, 1999). The cell model logic was adapted to the variable morphologies such that the original morphological organization of proximal, middle, distal dendrites was transformed to IML, MML, OML organization instead. The specific capacitance of the soma compartment was increased so that the mean input resistance measured at the soma was in the physiological range reported by Aradi and Holmes.

Basic morphological properties of the dentate interneurons such as total dendritic length and number of primary dendrites were obtained from experimental reconstructions reported in the literature (Bartos et al., 2001, Buckmaster et al., 1993, 2002, Freund and Buzsáki, 1998, Geiger et al., 1997, Lübke et al., 1998) and were used to create ball-and-stick morphological models with realistic total dendritic length, from which the total number of synapses was determined based on reported synaptic densities. However, detailed physiological data, including ion channel identities, was not readily available. In order to simplify the process of tuning the intrinsic biophysical properties of each cell, we adapted the Pinsky-Rinzel two-compartment model (Pinsky and Rinzel, 1994) to use an evolutionary optimization algorithm (Deb et al., 2002) to reproduce basic properties such as f-I curve, input resistance, membrane capacitance. Table 2 and Figure 8 summarize the intrinsic properties of each cell type, as well as the source used.

**Figure 8:**
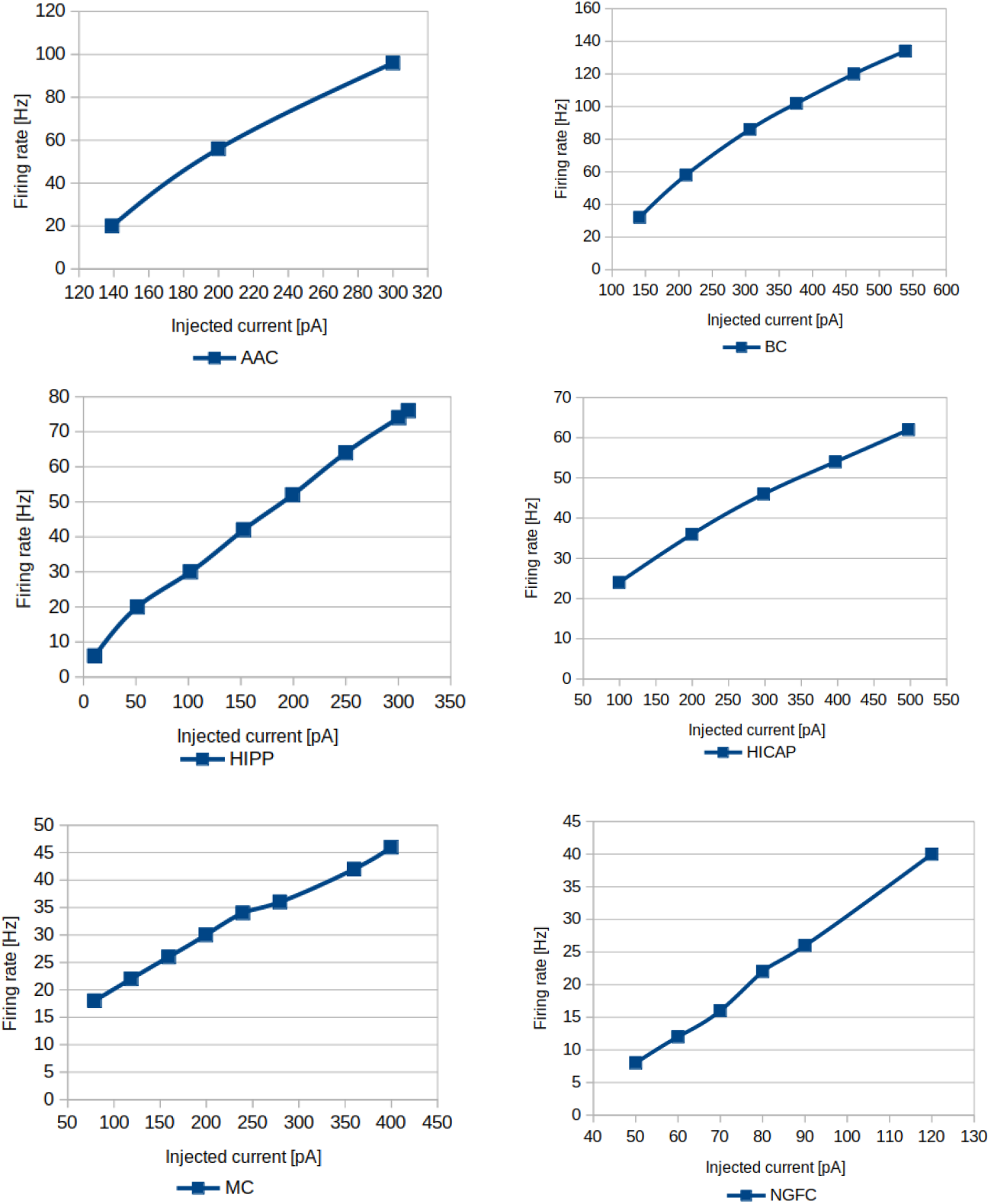
Frequency-current relationships of model neurons in the full-scale dentate gyrus model.

**Figure 9:**
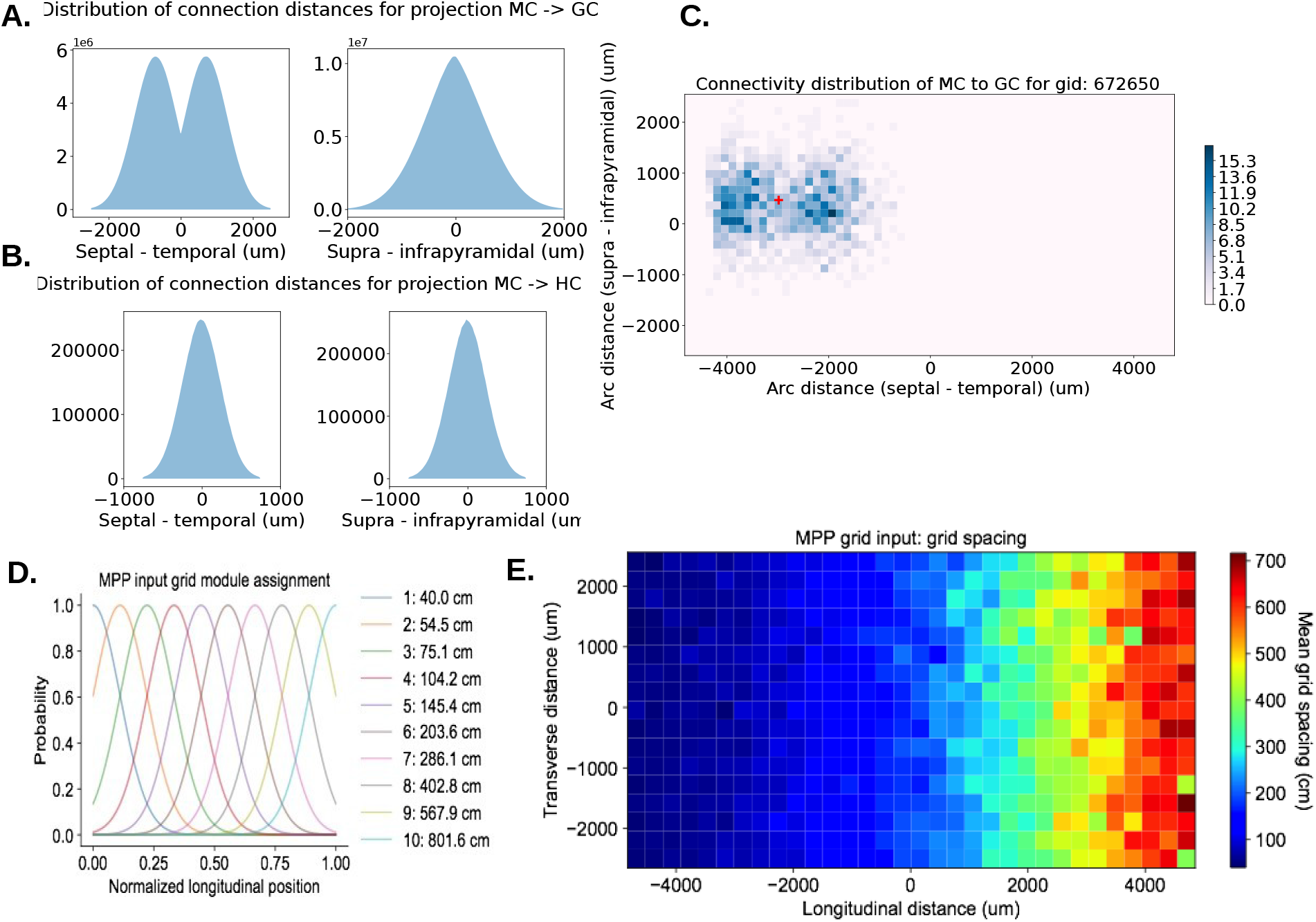
Topographical connectivity and input patterns of full-scale dentate gyrus model. A. Histogram of connectivity distances between mossy cells (MC) and granule cells (GC). Consistent with experimental reconstructions by Buckmaster et al. (Buckmaster et al., 1996), the mossy cell connectivity exhibits a characteristic bimodal distribution along the septotemporal axis. B. Histogram of connectivity distances between mossy cells (MC) and HIPP cells (HC). The mossy cell axon extent has been determined to be significantly shorter in the hilus than in the molecular layer, and the model connectivity reflects this accordingly. C. Plot of positions of presynaptic MC connected to a particular GC. Note that the majority of presynaptic MC tend to be located far away, reflecting the anatomical distribution of the MC axon. D, E. Topographical arrangement of the spacing of MEC grid cell inputs to the model.

**Table 2:** Intrinsic properties of model neurons in the full-scale model of the dentate gyrus.

| | Input resistance [$M\Omega$] | Memb. time constant [$ms$] | RMP [$mV$] | Spike threshold [$mV$] |
| --- | --- | --- | --- | --- |
| Mossy | 93 | 18 | -64 | -43 |
| Axo-Axonic | 175 | 3 | -65 | -45 |
| Basket | 150 | 2 | -62 | -46 |
| HIPP | 346 | 10 | -60 | -45 |
| HICAP | 171 | 18 | -60 | -43 |
| IS | 30 | 2 | -60 | -44 |
| NGF | 400 | 3 | -75 | -45 |
| MOPP | 400 | 3 | -75 | -45 |

Excitatory and inhibitory synapse counts for each cell type were estimated based on reported synaptic densities and total dendritic length. Synaptic locations for cell in the models were generated at intervals randomly sampled from Poisson distributions with parameters derived from the synapse density and compartment length at each segment of the cell morphology.

### 5.5 Synaptic Connectivity

Synaptic connectivity is summarized in Tables 5. Connectivity was estimated on the basis of experimentally determined numbers of connections and the population sizes of the pre-synaptic and post-synaptic cell types, based on a previous network and structural models (Dyhrfjeld-Johnsen et al., 2007, Schneider et al., 2012) and adjusted to account for the aforementioned changes in cell numbers and recent experimental evidence. Granule cells were previously estimated to make connections to 18 combined basket cells and axo-axonic cells, with no preferential targeting of either type. The previous 5:1 ratio was updated for the new prevalence of each type, so granule cells now contact basket cells and axo-axonic cells at 8:1 ratio. The previous estimate of 800 connections from mossy cells to HIPP and HICAP cells assumed lack of target preference and was updated using new prevalence ratios. Mossy cells connectivity was adjusted to account for differential axonal distributions in the hilus and the molecular layers, as well as for the additional CA3c backprojection and contralateral mossy cell inputs (see Table 5).

A recent study has found PV+ basket cell to HICAP cell connections at a similar level of connectivity as homogeneous PV+ basket cell connections (Savanthrapadian et al., 2014). As a result, this connection was added at ratio appropriate for the smaller prevalence of HICAP/CCK+ basket cells. Furthermore, the same study found no connections between HICAP and HIPP cells, but HIPP cells were found to contact other HIPP cells, and the connectivity structure of the model reflects these patterns accordingly. Neurogliaform and MOPP cells used similar connectivity patters, except inputs from HICAP and mossy cells to neurogliaform cells are not present, as the latter do not have soma or dendrites in the inner molecular layer (Armstrong et al., 2011).

Synaptic connections were selected by drawing samples from a Gaussian distribution, based on the longitudinal and transverse axonal distributions for the presynaptic cell type and the arc distances between the somata along these axes. The longitudinal and transverse connection extents for each cell type are summarized in Table 3, which are based on experimental measurements (except for MOPP cells, which is based on neurogliaform cell data). The connection probability distributions were centered at the septotemporal position of the soma, except for mossy cells, where the distributions were centered at distance 750 *μm* from the soma, according to the bimodal distribution observed in axonal reconstructions (Buckmaster et al., 1996). Where data was absent about the transverse axonal extent of a given cell type, it was assumed the same as for the longitudinal extent. The directional extent was set to represent three standard deviations for the Gaussian distribution, which assumes that approximately 97% of the axon lies within the reported extent, accounting for some axon collaterals being severed in the experimental slicing procedure.

**Table 3:** Longitudinal and transverse connection extents for all cell and input types in the full-scale model of the dentate gyrus. All distances are in units of micrometers.

|  | Hilus |  | GCL |  | IML |  | MML |  | OML |  |
| --- | --- | --- | --- | --- | --- | --- | --- | --- | --- | --- |
| Cell or input type | S-T | M-L | S-T | M-L | S-T | M-L | S-T | M-L | S-T | M-L |
| Granule | 900 | 900 | 250 | 250 |  |  |  |  |  |  |
| Mossy | 5,000 | 4,000 | 1,500 | 1,500 |  |  |  |  |  |  |
| Axo-Axonic | 2,000 | 1,000 | 2,000 | 1,000 | 600 | 300 | 400 | 200 | 20 | 10 |
| Basket | 600 | 300 | 1,700 | 1,000 | 1,200 | 800 | 100 | 100 |  |  |
| HIPP | 400 | 400 | 400 | 400 | 1,200 | 1,200 | 2,000 | 2,000 | 4,000 | 4,000 |
| HICAP | 2,600 | 2,600 | 1,300 | 1,300 | 2,600 | 2,600 | 1,500 | 1,500 | 500 | 500 |
| IS | 3,000 | 3,000 |  |  |  |  |  |  |  |  |
| MOPP | 400 | 400 | 200 | 200 | 1,250 | 1,250 | 2,000 | 2,000 | 800 | 800 |
| NGFC |  |  |  |  | 50 | 50 | 500 | 500 | 2,000 | 2,000 |
| CL Mossy | 1,500 | 1,500 |  |  | 5,000 | 4,000 |  |  |  |  |
| MPP |  |  |  |  |  |  | 1,500 | 3,000 |  |  |
| LPP |  |  |  |  |  |  |  |  | 1,500 | 3,000 |
| CA3c | 3,000 | 5,000 |  |  |  |  |  |  |  |  |

**Table 4:** Mean firing rates of identified neurons in the dentate gyrus during awake behavior.

| Neuron type | Mean firing rate [$Hz$] |
| --- | --- |
| Basket | 24.5 |
| Axo-Axonic | 24.1 |
| HIPP | 20.2 |
| HICAP | 4.4 |
| MOPP | 1.6 |
| NGF | 5.6 |

**Table 5:**
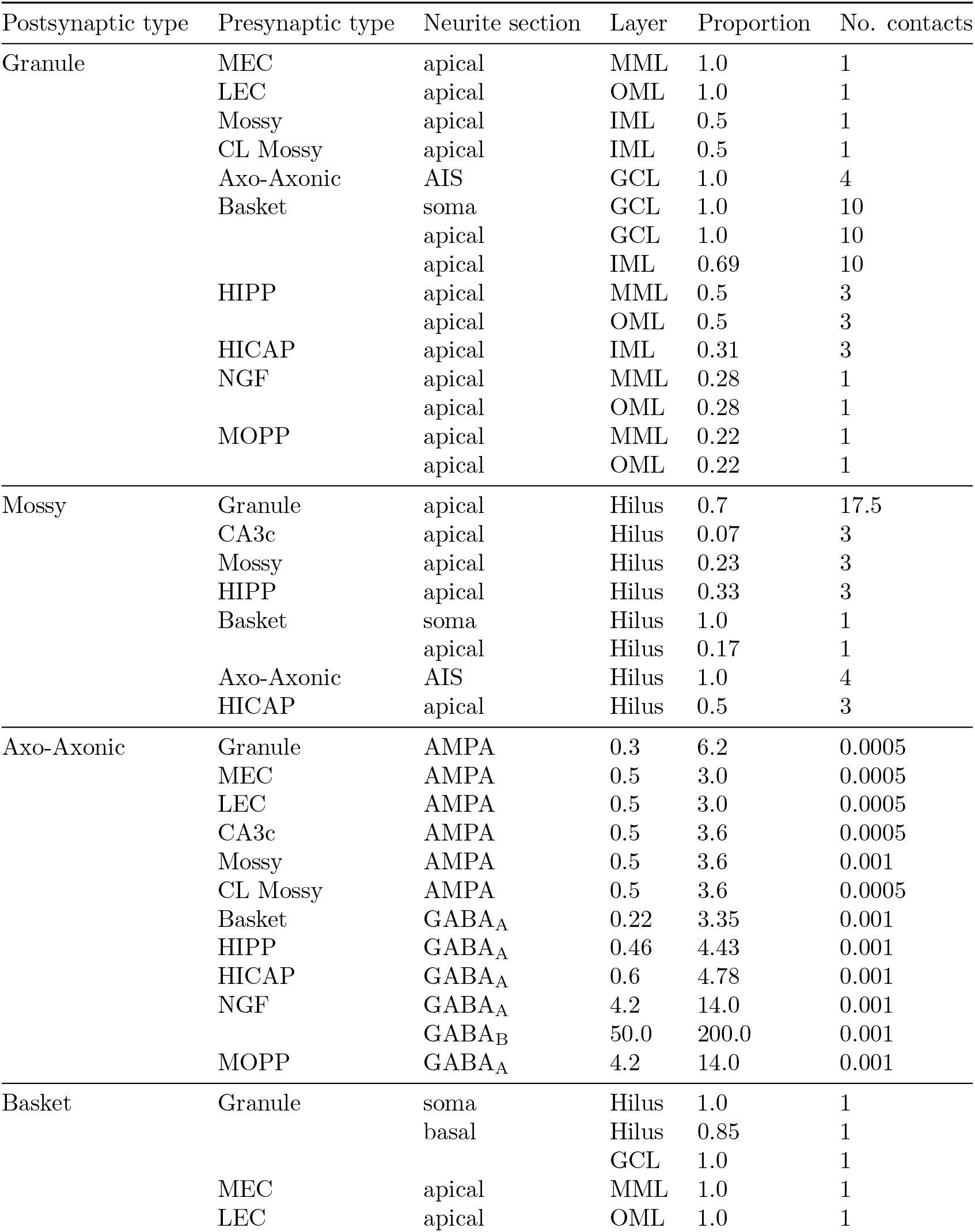
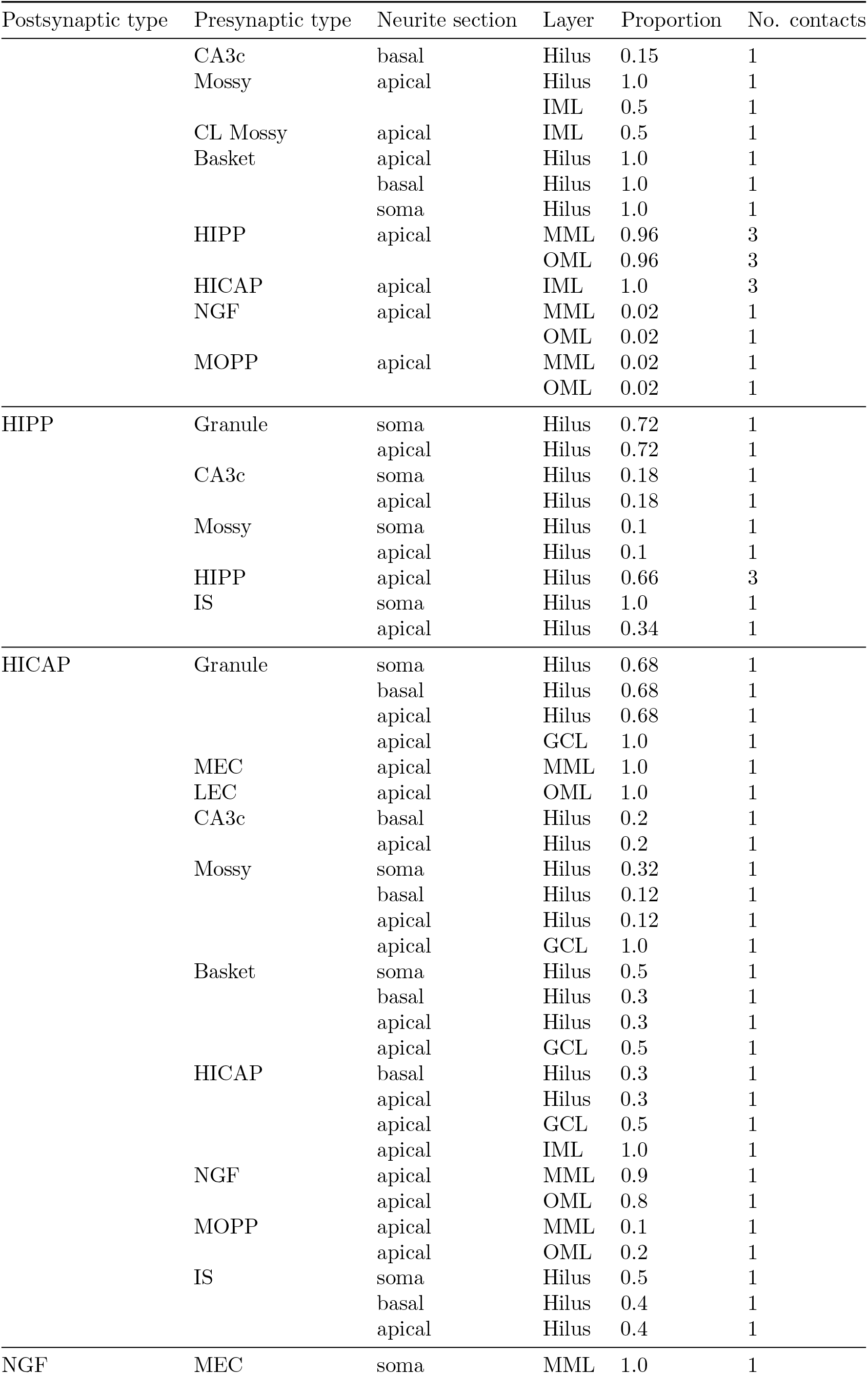
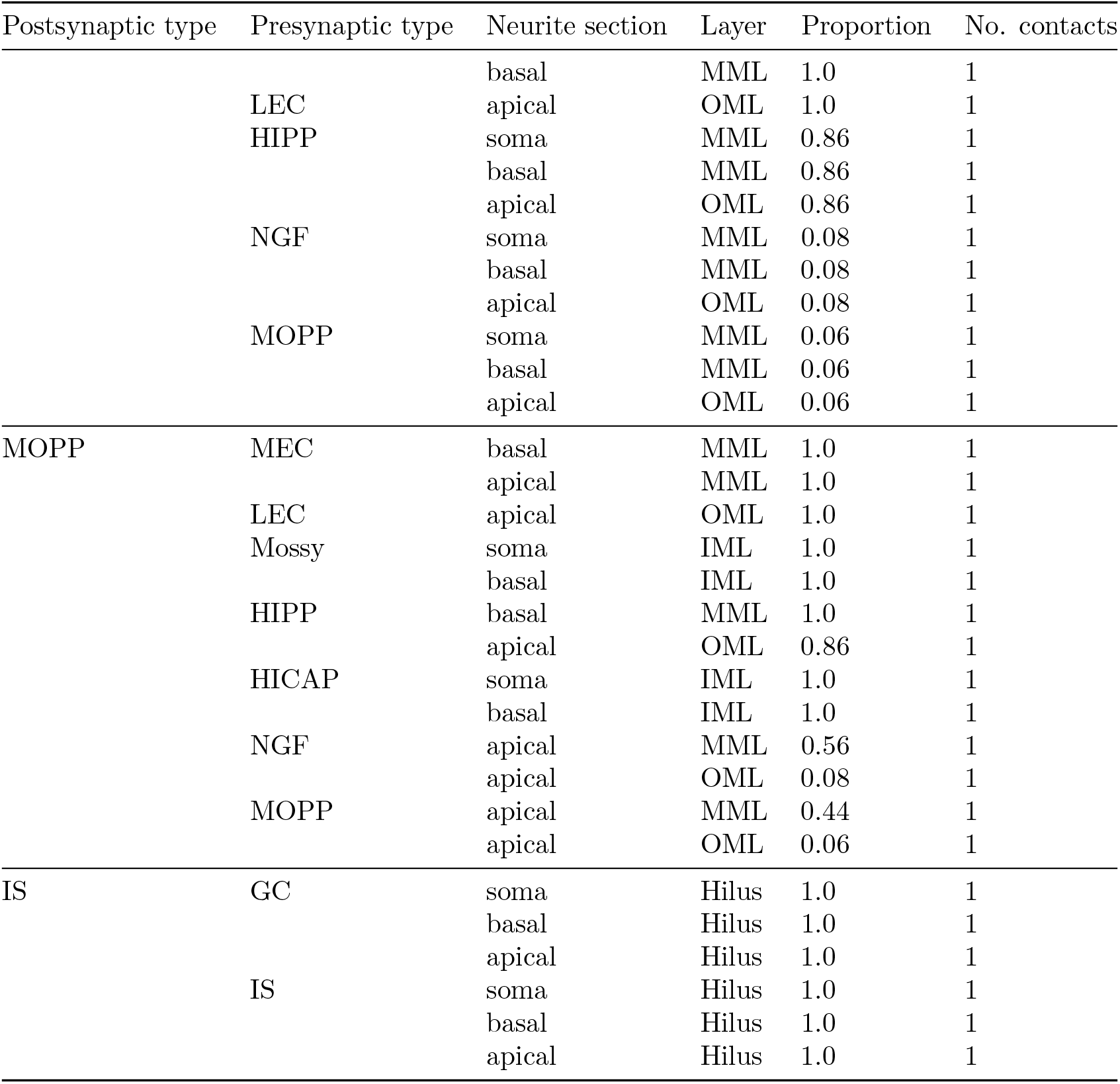
Connectivity parameters of full-scale model of the dentate gyrus.

The longitudinal and transverse distances between cells were approximated along the three-dimensional structure of the model dentate gyrus rather than computing exact arc lengths for each pair of cells. An interpolant for the longitudinal and transverse positions was created by sampling 1,000 points in the volume, measuring the arc distance to the edges of the volume by summing Euclidean distances along 1000 intermediate points, and fitting an Radial Basis Function interpolant with a Gaussian basis function as an approximate transformation between Euclidean and parametric surface coordinates.

The distance between a presynaptic source and a target cell was found by setting the layer parameter of the presynaptic position to the location of the target soma, interpolating the resulting arc distance, and adding the vertical distance between the layer positions of the source and target. The probability of connection was then sampled from a 2D Gaussian distribution based on the presynaptic axonal distributions. Synaptic pairs were then chosen based on the connection probabilities and number of connections for the given presynaptic cell type.

### 5.6 Receptor types and synapses

The maximum conductance, rise time and decay time constants for synaptic connections are summarized in Table 6. Maximum conductance was calibrated by measuring the EPSC or IPSC generated by the activation of a single synapse and adjusting the conductance to match available experimental data (Armstrong et al., 2011, Hashimotodani et al., 2017, Hefft and Jonas, 2005, Kraushaar and Jonas, 2000, Lysetskiy et al., 2005,

**Table 6:**
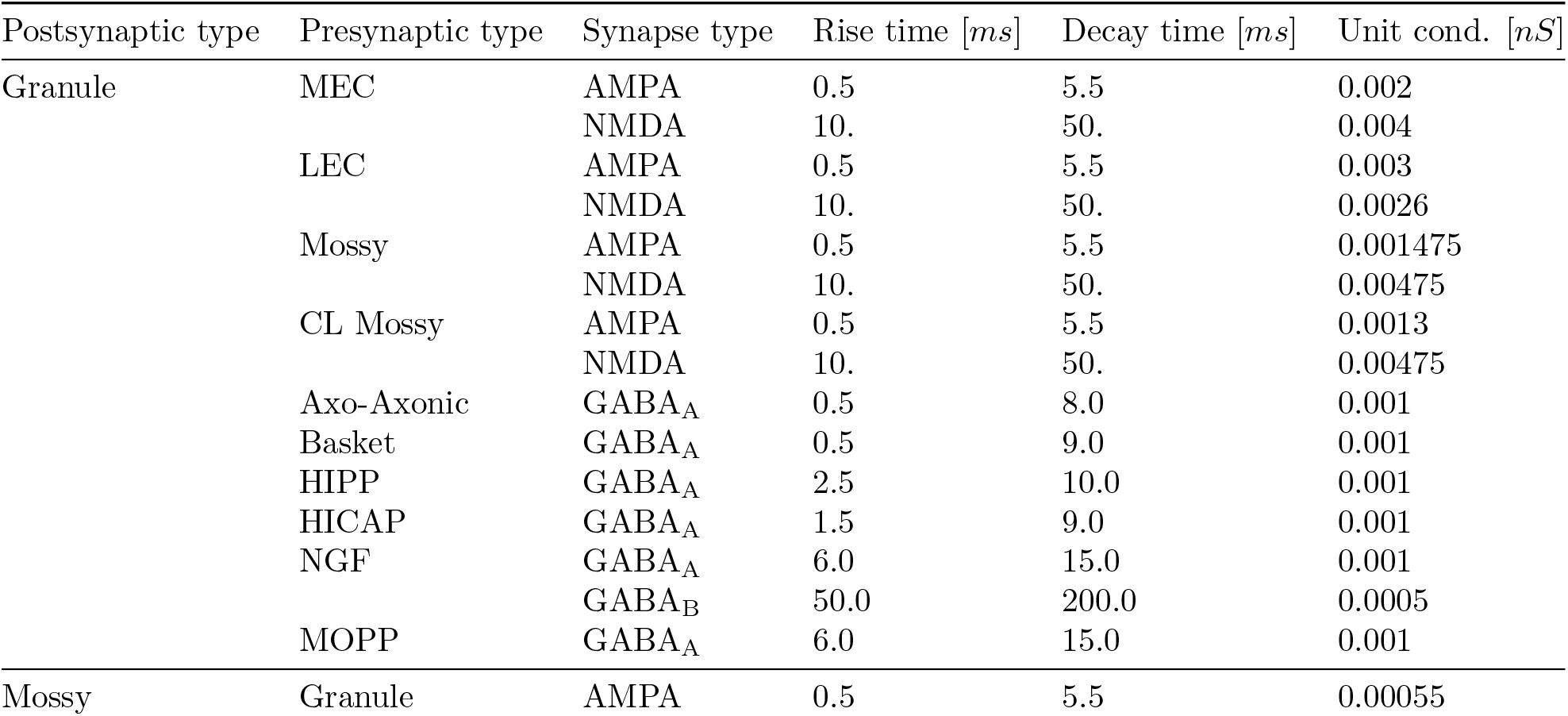
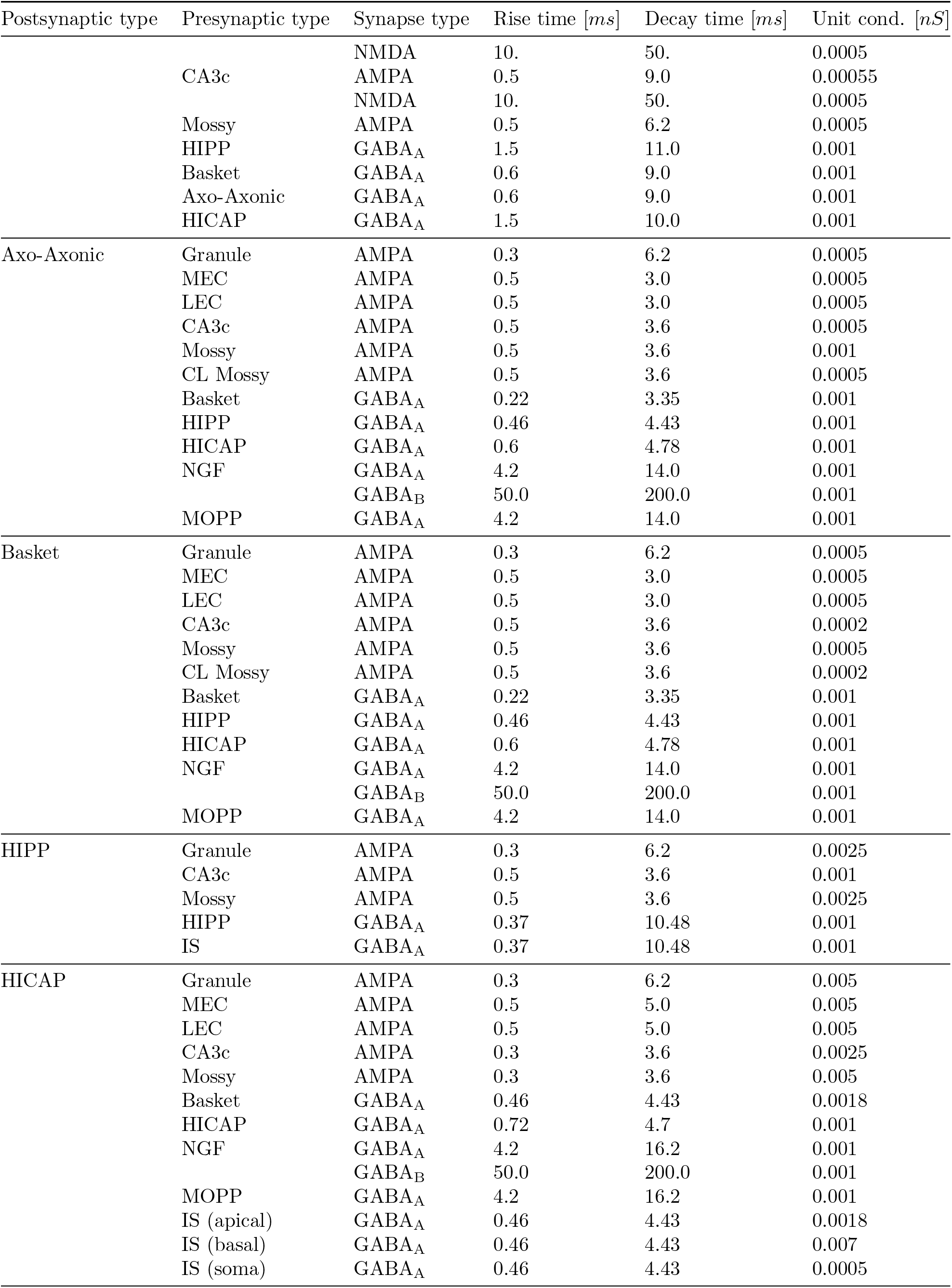
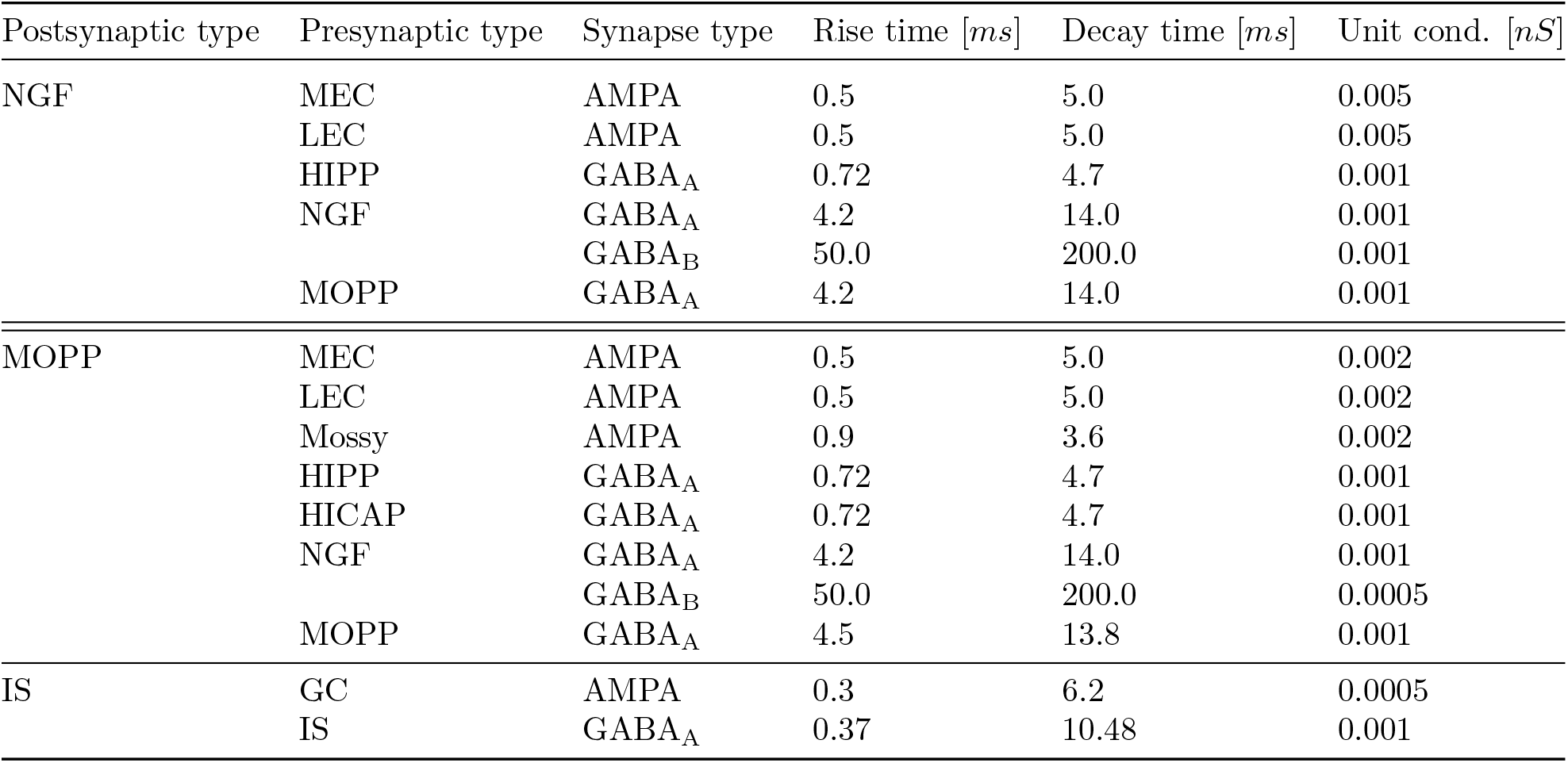
Synaptic parameters of full-scale model of the dentate gyrus.

Savanthrapadian et al., 2014). The synaptic weights of inhibitory interneurons were optimized via Network Clamp such that each cell type would fire at the mean firing rate recorded in-vivo during awake behavior (Table 4). Synaptic weights for connections thought to be shaped by plasticity processes (EC—granule cell, granule cell—mossy cell, CA3c—mossy cell) were randomly sampled from log-normal distributions with parameters derived from an EM spine size study by Trommald and Hulleberg (Trommald and Hulleberg, 1997). The weights of synaptic connections where associative plasticity is thought to be occurring (EC— granule cell, CA3—mossy cell) were additionally modulated by a supervised learning procedure based on the LSMR (Fong and Saunders, 2011) and L-BFGS (Liu and Nocedal, 1989) optimization algorithms were used to calculate the change of weights necessary to produce a place field at a predetermined random location in the simulated arena (Figure 10).

**Figure 10:**
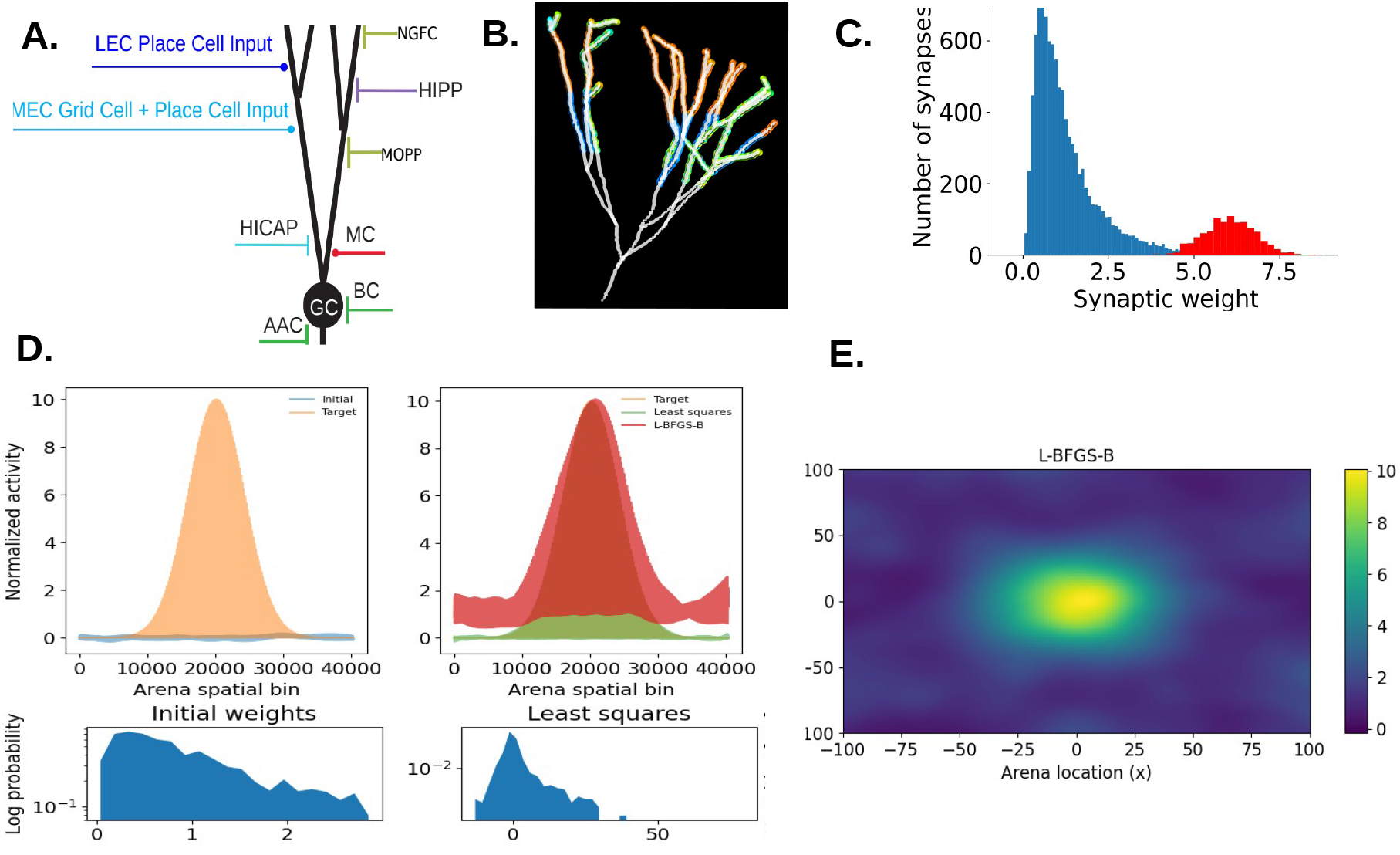
Cell-specific Distribution of Synaptic Properties. A. Schematic indicating the major source of excitatory and inhibitory inputs to dentate granule cells in the full-scale model of the dentate gyrus. B. Visualization of a synthetic granule cell morphology with color-coded excitatory synapses connected to MEC and LEC sources. C. Conceptual illustration of the synaptic weight distribution used in the model granule and mossy cells. Excitatory synapses are assigned weights sampled from a log-normal distribution in order to mimic the distributions resulting from long-term plasticity processes (Bromer et al., 2018) (blue histogram). Additionally, a subset of neurons are assigned place fields and a supervised learning procedure is used to mimic the potentiation of weights induced by associative learning. D1-4. Receptive field learning procedure. D1. Initial and target firing rate map in the simulated arena. The initial firing rate is calculated as the dot product of the initial synaptic weights with the firing rates of the inputs active at each bin of the simulated arena. The target firing rate is calculated as a Gaussian distribution centered on the target spatial receptive field. D2. Firing rates computed through synaptic weights obtained via least squares and local optimization procedures. D3. Initial distribution of synaptic weights. D4. Final distribution of weights computed by the learning algorithm. E. Firing rate map calculated through the final distribution of synaptic weights.

**Figure 11:**
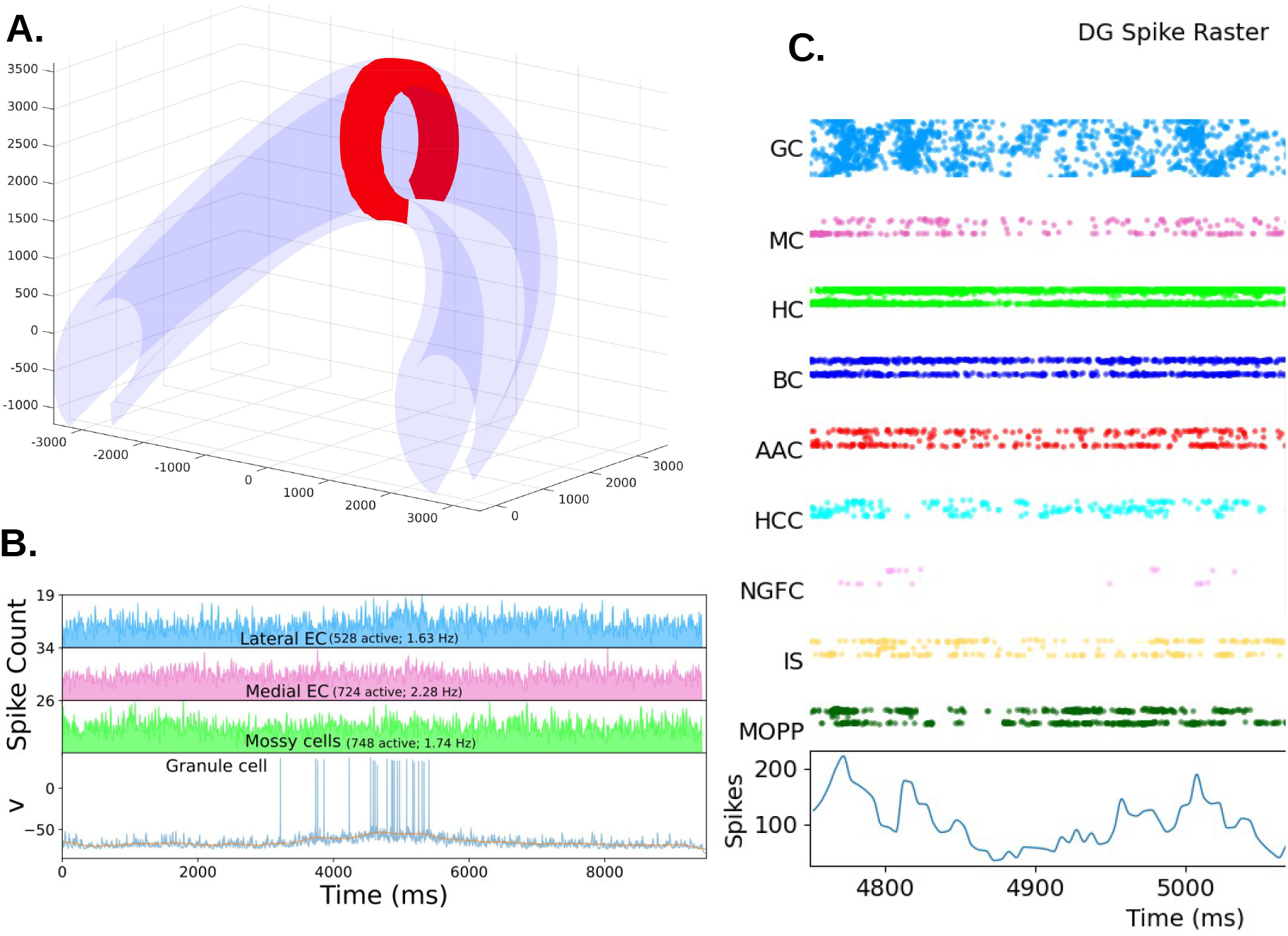
Results of synaptic parameter tuning of dentate gyrus circuit via Network Clamp / Microcircuit Clamp. A. A virtual slice can be extracted from the dentate gyrus model by retrieving all cells whose somata are within a geometric region of interest. B. An example Network Clamp simulation of a single granule cell with synaptic weights tuned to produce a spatial receptive field at the center of the simulated arena. C. Spike raster of a 50,000 cell model slice simulation.

### 5.7 Simulation Results

Our large-scale modeling framework enabled the declaration and definition of the previously described neuron, synapse and connection type-specific properties critical for testing the stability and robustness of our network. The full-scale dentate gyrus model contains 1,000,000 granule cells, each receiving 5000–10,000 excitatory synaptic inputs, including spatially modulated inputs from entorhinal cortex, long-range projections from mossy cells from the contralateral hemisphere, and projections to mossy cells and hilar interneurons from the CA3 region. The granule cell synapses are endowed with voltage-dependent NMDA receptors, and non-uniform synaptic distributions that are intended to mimic the results of the processes of long-term synaptic plasticity.

We initiated activity in the network via input spike patterns that represented the activity of entorhinal grid cells, and place cells, and CA3 place cells during locomotion. The input spikes are generated by simulating movement along a linear trajectory in a 2-D environment, and sampling spike times from an exponential distribution according to the expected firing rates of the input cells whose spatial receptive fields overlap with the positions along the trajectory. Additionally, the input spike patterns were theta phase modulated according to available experimental data (Sanchez-Aguilera et al., 2021). After an initial equilibration period where all inputs were gradually allowed to reach their target firing rates, rhythmic activity in the gamma range reliably emerged in the baseline network.

The network displayed stable and robust average firing activity, with low variability within the duration of the simulated trajectory path. The granule cells firing was ultra-sparse ( 8% active along the simulated trajectory), yet high rates were reached within their receptive fields (Figure 12). Overall, the average activity of various neuron types was-well aligned with their expected characteristics: the PV+ perisomatic cells (basket and axo-axonic cells) fired at the highest frequency, followed by the HIPP cells; granule cells displayed sparse and selective activity; mossy cells displayed selective activity; and the remaining interneurons fired within their experimentally observed low to moderate firing rates. The baseline network this constructed can be easily adapted to test different input configurations and hypotheses about different cell types and projections.

**Figure 12:**
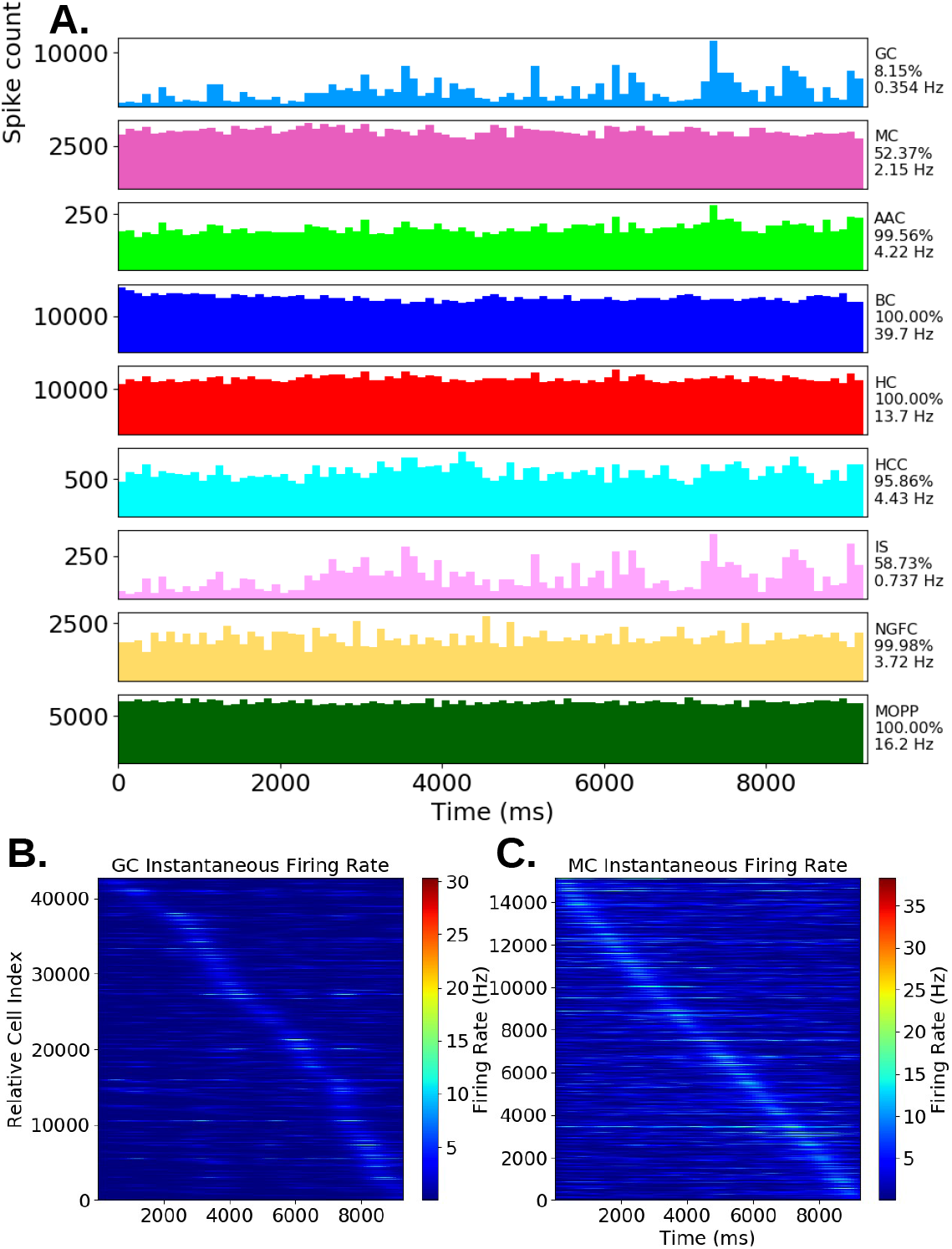
Results of simulation of full-scale dentate gyrus model. A. Spike histogram, fraction active, and mean firing rate of the biophysical neuron populations contained in the model. B. Instantaneous firing rate plots of granule cells and mossy cells in the model, highlighting the selective activity of those two cell types.

## 6 Discussion

Taken together, the results obtained through DG model simulation and optimization indicate that the connectivity patterns of granule cells, mossy cells, and interneurons as represented in our model of the dentate gyrus result in network dynamics that are consistent with the long-hypothesized lateral inhibition mechanisms mediated via mossy cells and GABAergic interneurons. We find that that the inhibition recruited through feed-forward inputs, along with feedback inhibition recruited via granule cells, mossy cells and hilar interneurons provides sufficient conditions for highly sparse activity of granule cells, and thus the unique connectivity structure of dentate gyrus may serve to enforce sparsity and, by extension, pattern separation.

Furthermore, recognizing the long-standing debate in computational neuroscience regarding the role large-scale neural simulation will yield novel insight into cognitive phenomena merely by virtue of being large-scale, we highlight possible strategies that employ large-scale models to make predictions about behaviorally-relevant neurobiological phenomena.

Most biophysical circuit models of the mammalian brain have treated the target brain region as homogeneous, with networks that consist of populations of neurons with identical intrinsic properties and synaptic connectivity. Yet the existing evidence points to significant anatomical and functional differentiation of the hippocampal microcircuitry along the three principal axes: the topographical projection patterns of EC to the hippocampus have been linked with dorso-ventral and proximo-distal gradients of place field properties (Henriksen et al., 2010, Jung et al., 1994), and significant functional differences between pyramidal neurons along the deep-superficial radial axis in CA1 (Slomianka et al., 2011).

In the cortex, there are several lines of evidence that hierarchical gradients of microcircuit properties determine large-scale specialization of cortical function (Huntenburg et al., 2018), and that one possible mechanism for processing information on different temporal scale is regional gradients in local synaptic properties reflected by microanatomical measurements of dendritic spines on pyramidal neurons (Palomero-Gallagher and Zilles, 2019, Scholtens et al., 2014).

Large-scale modeling provides the ability to define gradients in intrinsic neuronal and synaptic properties, connection and input patterns, as well as to define long-distance projections that otherwise would not be possible. Coupled with an efficient software infrastructure for the rapid generation and simulations of models of hypotheses about neuronal structure and function, large-scale modeling can provide detailed understanding how the interactions between gradients in local properties and network organization of brain regions shapes overall information processing in the brain.

## 7 Conclusion

We have presented key aspects of our scalable computational infrastructure for neuronal network modeling, which has been applied to construct, simulate, and analyze key information processing properties of a full-scale model of the hippocampal dentate gyrus (DG), and determine parameters that allow the model to generate sparse and spatially selective population activity that fits well with in-vivo experimental data. Our parametric computational representation of the reconstructed hippocampal volume allows the definition of unique heterogeneous morphological and biophysical properties of each model neuron, and allows highly detailed representation of topographic connectivity and afferent excitation provided from models of spatially-modulated inputs from entorhinal cortex, CA3c, and mossy cells in the contralateral hemisphere. In order to tune the firing rates of different cell classes, we have utilized our Network Clamp and Microcircuit clamp approaches, which involves the extraction of single neurons or virtual slice of the full-scale network and performing simulations and parameter optimization over the extracted cells contained, while providing idealized spike trains in place of the neurons outside of the volume. This work represents major progress towards solving the challenges of rapidly and efficiently constructing and evaluating large-scale biophysically-detailed network models with the goal of evaluating hypotheses about brain function that would be experimentally intractable.

## 8 Software Availability

The NeuroH5 library is available on the GitHub code sharing website at https://github.com/soltesz-lab/ neuroh5. Our prototype modeling framework and dentate gyrus model implementation is available at https://github.com/soltesz-lab/dentate.

## 9 Acknowledgments

The authors wish to extend thanks to Michael Hines and Pramod Kumbhar for their technical assistance with the NEURON and CoreNEURON simulators, and to Gerd Heber for technical assistance with the HDF5 format. Simulations were performed on the Frontera supercomputer at the Texas Advanced Supercomputing Center, University of Texas. Computing time was provided by NSF LRAC allocation FTA-Soltesz on Frontera. This work was supported by NIH (BRAIN U19 award NS104590).

